# The role of land use types and water chemical properties in structuring the microbiomes of a connected lake system

**DOI:** 10.1101/401299

**Authors:** Sophi Marmen, Lior Blank, Ashraf Al-Ashhab, Assaf Malik, Lars Ganzert, Maya Lalzar, Hans-Peter Grossart, Daniel Sher

## Abstract

Lakes and other freshwater bodies are intimately connected to the surrounding land, yet to what extent land-use affects the quality of freshwater and the microbial communities living in various freshwater environments is largely unknown. We address this question through an analysis of the land use surrounding 46 inter-connected lakes located within 7 different drainage basins in northern Germany, and the microbiomes of these lakes during early summer. Lake microbiome structure was not determined by the specific drainage basin or by basin size, and bacterial distribution did not seem to be limited by distance. Instead, land use within the drainage basin could predict, to some extent, NO_2_+NO_3_ concentrations in the water, which (together with temperature, *chlorophyll a* and total phosphorus) affected water microbiome structure. Land use directly surrounding the water bodies, however, had little observable effects on water quality or the microbiome. Several microbial lineages, including environmentally important Cyanobacteria and Verrucomicrobia, were differentially partitioned between the lakes. As the amount of available data on land use (e.g. from remote sensing) increases, identifying relationships between land use, aquatic microbial communities and their effect on water quality will be important to better manage freshwater resources worldwide, e.g. by systemically identifying water bodies prone to ecological changes or the presence of harmful organisms.

## 1. Introduction

The world’s population growth is maintained by a suite of human enterprises such as agriculture, industry, fishing and international commerce (Vitousek, *et al*., 1997). The use of land for such enterprises has greatly altered the way ecosystems interact with the surrounding land, atmosphere and aquatic systems (Vitousek, *et al*., 1997). Freshwater ecosystems, in particular, may be highly sensitive to human impact, mainly through eutrophication. The microbial communities (“microbiomes”) living in freshwater ecosystems are intimately connected with water quality. Microbes take up, utilize and recycle elements such as carbon, nitrogen, phosphorus and sulfur, and thus control and modulate local and global elemental cycles (Newton, *et al*., 2011). Some aquatic microorganisms are potential pathogens of animals or humans, or can produce secondary metabolites or toxins (Abdelzaher, *et al*., 2010, Cabral, 2010, Valério, *et al*., 2010). Thus, significant effort has been invested in elucidating the environmental drivers that determine the composition and function of freshwater microbiomes (reviewed by, e.g., Logue & Lindström, 2008, Williamson, *et al*., 2009, Newton, *et al*., 2011, Zeglin, 2015). Environmental drivers of microbial community structure may act at the local scale, where the physical-chemical characteristics of the water lead to a “species sorting” process which enrich for bacteria adapted to occupy that specific niche (Jones & McMahon, 2009, Langenheder & Székely, 2011, Van Rossum, *et al*., 2015). Such physical - chemical properties can include, for example, nutrient, organic carbon or metal concentrations, pH, temperature, salinity and light intensity (Crump, *et al*., 2004, Horner-Devine, *et al*., 2004, Yannarell & Triplett, 2005, Zeglin, 2015). The physical-chemical properties of water depend, in turn, at least partly on the input of nutrients, organic matter and other contaminants from the surrounding terrestrial ecosystem via the water shed (Chen, *et al*., 2018). Specifically, different types of land use surrounding the water bodies (e.g., forests, agricultural lands, urban lands, etc.) can strongly affect the quantity and quality of terrestrial input into water bodies (reviewed by (Solomon, *et al*., 2015).

Local processes can be affected by processes occurring at larger spatial and temporal scales. For example, physical connectivity between water bodies (e.g., within the same drainage basin) may lead to bacterial transfer between environments, a process termed “mass effect”, (e.g., Quinn, *et al*., 1997, Nelson, *et al*., 2009, Crump, *et al*., 2012, Lindström & Langenheder, 2012, Logue, *et al*., 2012). In very strongly connected habitats, dispersal rates may be so high that they lead to ecosystem homogenization (Leibold & Norberg, 2004). As the spatial scales increase, geographic distances and physical barriers may lead to limitations on bacterial dispersal, resulting in “distance decay” patterns, whereby differences between microbial communities increase with distance (Van der Gucht, *et al*., 2007, Lear, *et al*., 2013, Niño-García, *et al*., 2016, Marcelino, *et al*., 2017, Logares, *et al*., 2018). As the various processes affecting freshwater microbial ecosystems are identified and characterized, many questions remain: to what extent are local conditions within water bodies affected by those in the surrounding terrestrial system? What is the relative importance of local conditions in determining microbial community structure? What is the role of mass transport of microbes from upstream sites, and how does this differ between freshwater systems? Are all microbial taxa affected in the same manner by these processes? While many of these questions have been addressed in ecosystems ranging from highly impacted agricultural lands (e.g. Griffin, *et al*., 2017) to pristine Antarctic lakes (Logares, *et al*., 2018), few studies have aimed to integrate the effects of hydrology and water chemistry on lake biology in the context of land-use within their respective catchment basins (e.g. Catherine, *et al*., 2016). Answering some of these questions is critical as we aim to understand how changes in the environment, including land use, affect aquatic ecosystems.

In this study, we investigated the microbiomes of 46 German lakes, located in a network of 7 drainage basins connected through streams. The region encompassing these lakes and basins comprises a tapestry of different natural and artificial land parcels (e.g., forests, pastures, agricultural lands, water bodies, towns and villages, Figure 1). Our major goal was to understand what shapes microbial population in such a system of connected natural water bodies, located within relatively short distances and surrounded by a variety of different land-use types. We hypothesize that: 1) land use directly affects water chemistry; 2) water chemistry affects bacterial composition by habitat filtering of bacteria adapted to specific water properties; 3) bacterial populations gradually change as the water moves downwards within the connected lakes and streams; 4) there may be also a direct effect of land use types on the water microbiome via direct input of live bacteria.

**Figure 1.**
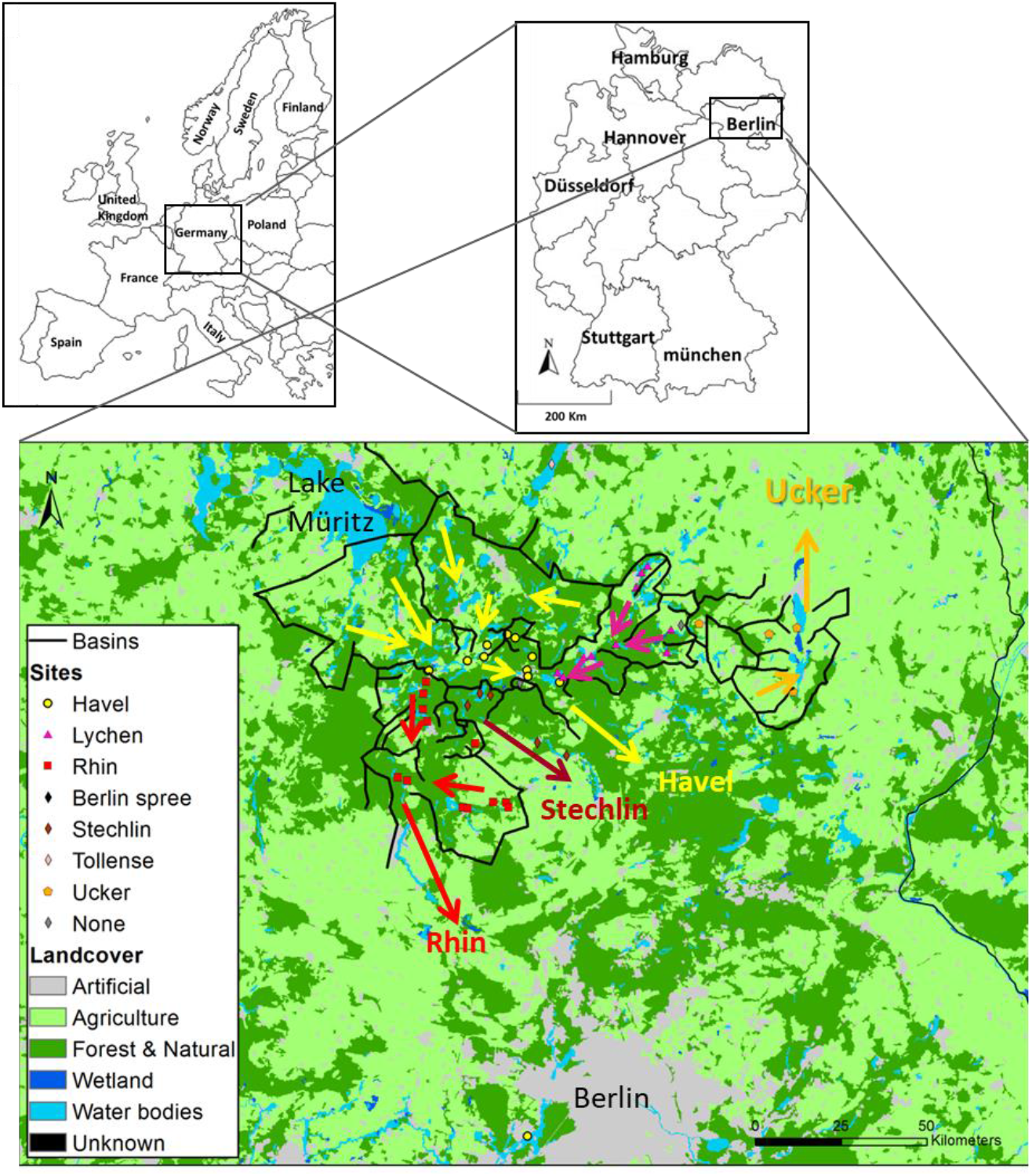
Land use and drainage basins in the sampling area. A total of 46 samples were collected from lakes and a fish-pond in the Brandenburg region of northern Germany. Five types of land use: urban areas, agriculture (further divided for GLM analysis to heterogeneous agriculture, pasture lands and arable lands), forests, wetlands and water bodies, were mapped using GIS, with black lines superimposed indicating drainage basins. The arrows indicates the dominant flow direction within each basin. A map with the identities of each sampling location is presented in Supplementary Figure 1, and the detailed information on each sampling location can be found in Supplementary Tables 4–6.

## 2. Materials and methods

### 2.1 Study region and delineation of drainage basins

The sampling area lies within the region of Brandenburg, in northern Germany (Figure 1, Supplementary Figure 1, Supplementary text). The area is relatively flat (18-89 meters above sea level) and contains hundreds of lakes, ponds and depressions that were formed by glacial action during the Pleistocene. Three main basins drain this region, which are (from west to east) the Rhin (which flows north), the Havel and the Ucker, the two latter flowing south in the region of sampling. These three main basins can be subdivided into smaller basins. The drainage basin of each lake was defined as the line connecting the highest points from which the surface water flow enters each of the lakes, including all of the upstream lakes. We note that groundwater flow may be significant in many lakes, and often does not flow as surface water does in accordance with topographic basins (Winter, *et al*., 2003). We also assume that the water is fully mixed horizontally within each separate water body, and that our sampling represents the epilimnion if the lakes are stratified. A detailed description of the sampling region, methods of delineating the basins, identification of the canals connecting the basins and GIS layer analysis, can be found in the supplementary information.

### 2.2. Sampling and analysis of environmental data

Each location was sampled once from the edge of the water body, as due to the range of water bodies sampled and the inaccessibility to some by boat sampling from the deepest point was not feasible. During sampling, dissolved oxygen, temperature pH and turbidity were measured using a YSI multi probe (YSI 6920 V2-2; YSI Inc). At all locations, 5 liters of surface water were collected using plastic jerry-cans that were washed with double distilled water then with 70% ethanol and air dried prior to sampling. Care was taken not to raise any sediment. Samples were brought to the lab and filtered within two hours. For *Chlorophyll a (chl* a) measurement, water was filtrated on GF/C filters (Whatman glass fiber, 25 mm, nominal pore size 1.2 μm) and placed into 1.5 mL sterile Eppendorf tubes. The water was filtered until the filters were clogged, and the volume was recorded (50-800 mL). *Chlorophyll a* was extracted in absolute methanol for 12 h at room temperature in the dark, and the concentration was determined spectrophotometrically according to (Ritchie, 2008). For DNA extraction, water was filtered using a vacuum pump on GF/F filters (Whatman glass fiber, 25 mm, nominal pore size 0.7 μm). Preliminary experiments using a culture of the marine Pico-Cyanobacterium *Prochlorococcus* MED4 (~0.5 μm in diameter) showed that with the vacuum pumps we used approximately 99.9% of the cells of this size were retained on a GF/F membrane. The filters were placed into 1.5 mL sterile Eppendorf tubes with 1 mL of storage buffer (40 mM EDTA, 50 mM Tris-HCl, 0.75 M Sucrose) and stored at −80°C until analyzed. The filtrate from the GF/F filters was collected for measurement of dissolved nutrients, and unfiltered water was used for total phosphorus measurements according to (Grasshoff & Kremling, 1983, Grasshoff, *et al*., 2009) DNA extraction was performed using bead-beating followed by robotic extraction using a QiaCube robot, the gene encoding the 16S subunit of the ribosomal RNA was amplified using primers targeting the V3-V4 region, and the amplicons were sequenced using the 2×250 base pair format on a MiSeq flow cell (V3 chemistry). More details on the DNA extraction and 16S amplification procedures are found in the supplementary methods section.

### 2.3 Generalized Linear Model Analysis (GLM)

We used multivariable regression model in the framework of GLMs to relate the six land use types to the measured water chemical parameters. Multi-model inference based on the Akaike Information Criterion (AIC) was used to rank the importance of variables (Burnham & Anderson, 2003, Blank & Blaustein, 2014). The package ‘‘glmulti’’ was used to facilitate multi-model inference based on every possible first-order combination of the predictors in each scale (Calcagno & de Mazancourt, 2010). The coefficients associated with each variable and their relative importance were assessed using a multi-model average. Parameters were estimated by averaging across models, weighted by the probability that their originating models are the best in the set. Hence, the relative importance of a parameter is not based on *P*-values, but rather on its occurrence in the models that are best supported by the data. The importance value for a particular predictor is equal to the sum of the weights for the models in which the variable appeared. Thus, a variable that shows up in many of the models with large weights received a high importance value.

### 2.4 16S Sequence analysis and statistical tests

Sequences received from the sequencing facility were quality filtered to remove PhiX DNA, unmerged reads and low quality sequences, followed by classification into different Operational Taxonomic Units (OTUs) based on 97% sequence similarity threshold using the “pick_de_novo_otus.py” command in Qiime (Caporaso, *et al*., 2010). OTUs with fewer than 5 reads across all samples were removed, resulting in 35,775 – 75,704 reads per sample. Alpha diversity (Shannon index) was calculated using the ‘’diversity’’ function, and Bray– Curtis dissimilarity by ‘’vegdist’’ function. Non-metric multidimensional scaling analysis (nMDS) was carried out in R with ‘’metaMDS’’ function and a Bray-Curtis matrix. In order to relate environmental factors to specific OTUs we performed variation partitioning (“varpart” function in R) followed by canonical correspondence analaysis (CCA) using the “cca” and “anova.cca” functions in the “vegan” R package, on subsamples of 25,000 reads for each location which were then log10 transformed. For additional information on these analyses, see the supplementary methods.

### 2.5 Permutation test of phyla clustering significance

In CCA ordinations, the location of each OTU within the ordination is related to the effect of the variables constraining the CCA on that OTU. We wanted to address a different question - whether a group of OTUs (e.g. a phylogenetic lineage) behave coherently with respect to the environmental characteristics. In other words, we do not ask, for example, whether “a cyanobacterial OTU” is related to the CCA axes (as that is the direct output of the CCA), but whether the cyanobacteria as a group behave similarly with respect to these axes. We therefore designed a permutation test to determine: (a) the clustering significance of the OTUs within each clade; (b) the significance of association with the tested environmental factors. We obtained the coordinates of eigenvectors (e.g. pH, temperature, etc.), and of all OTUs belonging to the different clades, in the CCA data. For each clade *p*, we calculated two scores: *d*_p_ and *d*_p,v_, where *v* is a given eigenvector. The score *d*_p_ equals the sum of Euclidean distances between all pairs of OTUs in *p*. The score *d*_p,v_ is the sum of scalar projections of all OTUs in *p*, on a given eigenvector v. Accordingly: (a) for the clustering test, *s*_p_ OTUs were randomly chosen from all tested clades, to obtain the score *d*_p=random_, where *s*_p_ is the count of OTUs in clade *p*. This calculation was repeated n times (*n*=1000), where the count of cases where *d*_p_ ≥ *d*_p=random_, divided by *n*, was defined as p-value; (b) for the association test, The score *d*_p,v_ was compared to the score *d*_p=random,v_ which was obtained from random coordinates normally distributed with the same x,y variance as the x,y variance as in clade *p*, and the same number of *s*_p_ datapoints. This calculation was repeated n times (*n*=100), where the count of cases with *d*_p.v_ ≥ *d*_p=random,v_ divided by *n*, was defined as *p*-value. Here, *p*-value < 0.025 or > 0.975 (after Benjamini-Hochberg correction for multipole hypothesis testing) indicates negative and positive association with the tested environmental factor.

## 3. Results

### 3.1 General characterization of the sampled water bodies

During the summer of 2015, we sampled 46 water bodies (lakes and one fish pond) within a relatively small area (~150 km by 100 km) of the Brandenburg/Mecklenburg lake district in northeastern Germany (Supplementary Excel file 1). The lakes ranged in trophic status from meso-oligotrophic (Lake Stechlin) to polytrophic (hyper-eutrophic), with total phosphorus concentrations between 0.009-0.589 mg/L and *chl* a concentrations ranging between 0.0005-0.320 mg/L. The water bodies belonged primarily to five major drainage basins (Figure 1, Supplementary Figure 1), and varied widely in their sizes and nutrient concentrations (Supplementary Data file 1). While the microbial diversity, expressed as the Shannon alpha-diversity index, between the basins was relatively similar (4.88-7, Figure 2A), clear differences were observed in community composition between the various lakes (Figure 2B). Comparing the microbiome composition between lakes (using the Bray-Curtis dissimilarity index) revealed that they can be broadly divided into two main groups: a cluster of sites with a high relative abundance of Cyanobacteria, and a second cluster with a high relative abundance of Actinobacteria, Proteobacteria and Bacteoidetes and low relative abundance of Cyanobacteria (Figure 2B).

**Figure 2.**
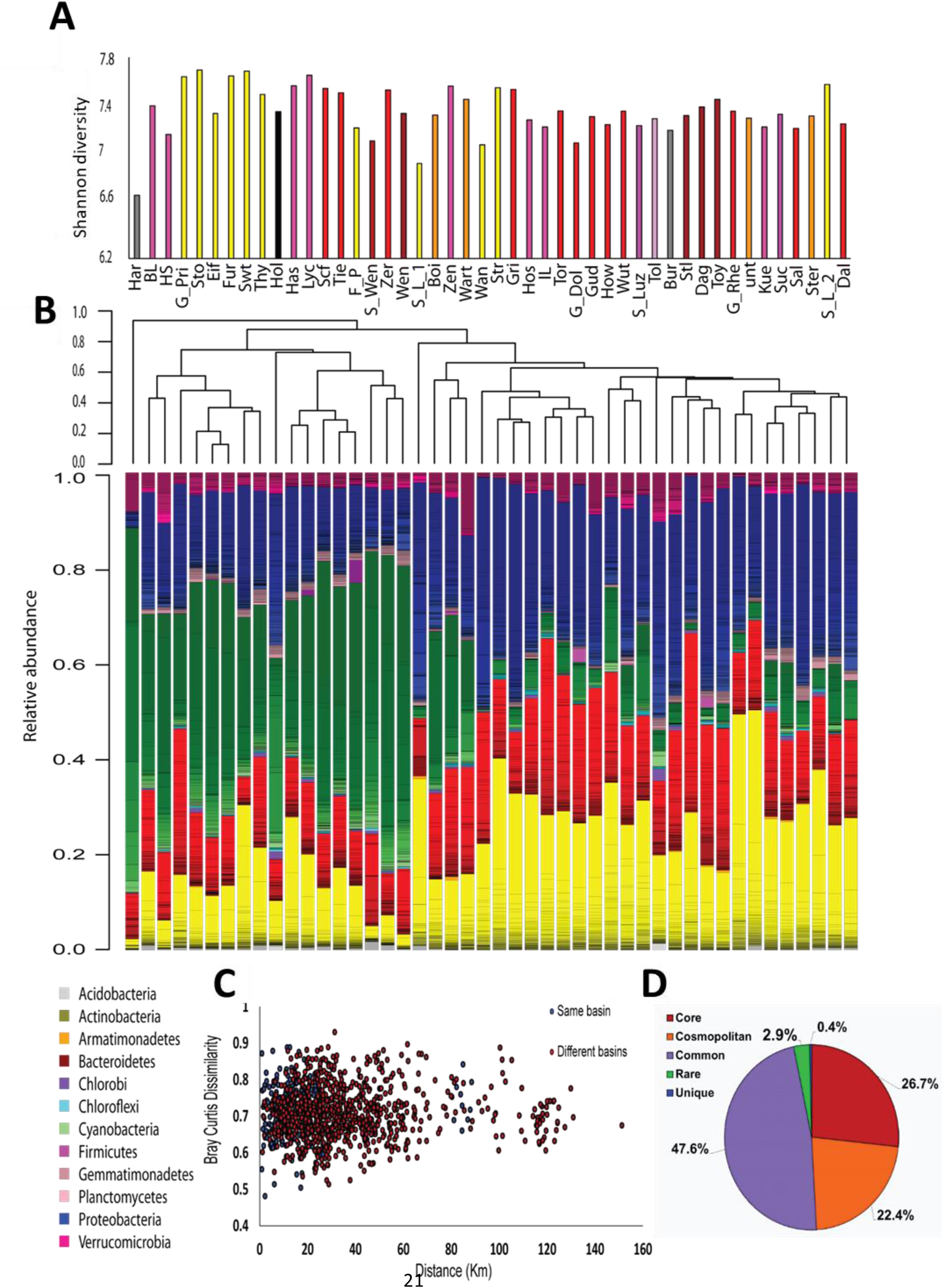
An overview of the microbial populations in the sampled lakes. **A) α-diversity of the microbial populations in the different lakes, measured by the Shannon index**. The columns are colored by drainage basins. No clear patterns are observed. **B) Community composition and community clustering based on the Bray-Curtis similarity index**; Colors represent different phyla, lines within the same color represent OTUs at the 97% identity cutoff. The sampling sites can be divided into two groups: one dominated by Cyanobacteria, and the other by Actinobacteria, Proteobacteria and Bacteroidetes. **C) No distance decay patterns are observed**. Each point represents the Bray-Curtis dissimilarity between locations within the same basin (blue) or locations found in two separate basins (red). Spearman’s r between geographic distance and Bray Curtis distance is 0.03, p=0.30. **D) Most of the OTUs are found in more than 50% of the sampling sites**. The diversity (γ diversity) of all sampling locations combined is presented, divided into five categories representing core OTUs (found in all sites), cosmopolitan OTUs (> 90% of the sites), common OTUs (> 50% of the sites), rare OTUs (< 10% of the sites), and unique OTUs (present at one site only).

Lakes are often broadly divided into trophic states, using parameters such as *chl* a or total phosphorus (TP) concentrations, and these trophic definitions are often used in water management. As shown in Supplementary Figure 2, the trophic status of the lakes based on *chl* a and total P concentrations was broadly in agreement with estimates based on multi-annual monitoring by the German federal states of Mecklenburg-Vorpommern and Brandenburg, although more of the lakes had an oligotrophic character at the time of sampling. This is likely because we sampled during early summer where high production rates of zooplankton can reduce phytoplankton abundance, leading to a “clear water phase” and masking the real, higher trophic status of lakes high in nutrients. A slight tendency was seen for lakes with higher *chl* a and total P concentrations to have a higher relative abundance of Cyanobacteria (Supplementary Figure 2) manifested also as an observable trend in the NMDS ordination (Supplementary Figure 3A).

Interestingly, lakes from the same drainage basins did not cluster together, despite relatively large differences between the drainage basins in terms of their size and land-use in their water shed (Figure 2B, Supplementary Figure 3B, Supplementary Data File 1). Neither was a clear correlation observed between size of the drainage basin of each water body and chemistry of the water (Supplementary Figure 4A-F) or lake microbiome (Supplementary Figure 4G). One major factor, that might determine to what extent ecological communities differ, is the geographical distance between them. This phenomenon, identified by a “distance decay” relationship in community similarity (Nekola & White, 1999, Bell, 2010), has been demonstrated for both macro- and micro-organisms at various scales of geographic distances (Lear, *et al*., 2013, Marcelino, *et al*., 2017). No distance decay relationships were observed in our samples, neither within a single basin nor between samples belonging to different basins (Figure 2C). The lack of distance-decay patterns was still observed when the three major basins, Lychen, Rhin and Havel, were divided into sub-basins where all of the water bodies were directly connected (Supplementary Figure 5). Moreover, examination of the total species diversity in our landscape (i.e. γ-diversity) showed that 27% of the OTUs are core OTUs that were present among all sites, and that additional 22% OTUs were cosmopolitan, defined as being observed in >90% of the sites (Figure 2D). Rare and site unique OTUs combined represented only 3.4% of all OTUs. These rare and unique OTUs were also numerically scarce, representing at most 2.6% of the total sequence reads. The core microbiome was dominated by Actinobacteria and Proteobacteria (43% and 39% of the OTUs respectively, Supplementary Figure 6A), and contained very few Cyanobacteria OTUs (1%). In contrast, Cyanobacteria represented the dominant phylum of all cosmopolitan OTUs (47%). Thus, it seems that differences between microbial communities are primarily due to changes in the relative contribution of OTUs rather than to the presence of unique or endemic OTUs in different regions.

### 3.2 Land use in the drainage basins affects water chemistry, and through it, the microbiome

We next asked whether specific land use types may affect environmental conditions within the water body (e.g. nutrients, temperature, etc., referred to here as “aquatic environmental conditions”, see supplementary information for the definition of the land-use types used). The effect of land use could be manifested at the level of the entire drainage basin (summing up the input of the entire draining into each water body), or, alternatively, at the level of local land use (adjacent to the water body). To test our hypotheses, we used General Linear Models (GLMs) to determine whether specific land use types could be associated with specific aquatic environmental conditions. On the basin scale, six land use types (agricultural area, pasture lands, arable lands, urban and forest areas and water bodies) explained a high proportion of the variability of nitrite and nitrate concentrations (NO_2_+NO_3_, R_N_^2^= 0.74, Figure 3, Supplementary Table 1), with the fraction of urban land being positively correlated to NO_2_+NO_3_ concentrations (Supplementary Table 1). When quantifying the independent contribution of each different land use separately, urban was the only significant variable explaining 74% of the variation in NO_2_+ NO_3_.

**Figure 3.**
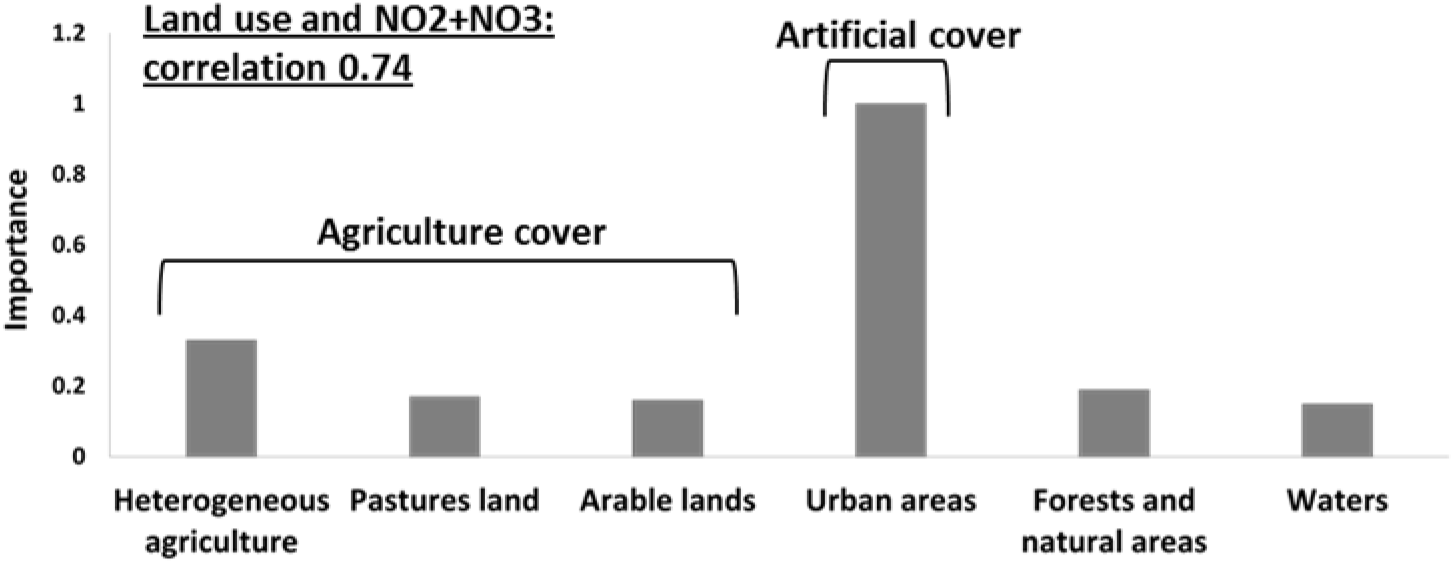
Spesific land use typy affects aquatic environmental parameters: General Linear Models (GLMs) analysis were performed and resulted in **Urban areas are positively related to NO_2_+NO_3_ concentrations**. A strong correlation (R^2^= 0.74) is observed between land uses and nitrite and nitrate concentrations, with the fraction of “urban areas” being the most important variable contributing to the correlation

We next used VPA to explore whether and to what extent basin-level and local-level land use types, as well as aquatic environmental conditions, explain the composition of the aquatic microbiome. VPA results pointed to aquatic environmental parameters as key parameters, in total explaining 15.5% of the variance in microbiome composition among all sites, 11% independent of land-use parameters (Figure 4). In contrast, land-use types (at both basin and local scale) had a very weak explanatory power for microbial community structure (Figure 4). Following VPA results, we tested measured aquatic environmental parameters as explanatory model for community composition using CCA followed by CCA permutation test (Supplementary Table 4, Figure 5). The aquatic environmental parameter model explained 24.2% of the inertia in CCA and was significant (*P*<0.05). Similar to the NMDS (unconstrained) ordination (Supplementary Figure 3A), the CCA did not show clear separation of sampling sites by basin (Figure 5b). Four out of six aquatic environmental parameters in our model were significant: total phosphorus (TP, *P* < 0.01) and *Chl a* (*P* < 0.05) concentration were positively associated with CCA1 (eigenvector for CCA1 and CCA2 were 0.89, −0.12 for *Chl a* and 0.45, −0.16 for TP), NO_2_+NO_3_ (*P* < 0.05) was positively associated with CCA2 (eigenvector 0.30, 0.58) and water temperature (*P* < 0.01) was negatively associated with CCA2 (eigenvector 0.08, −0.84) (Figure 5a, Supplementary Table 4). Notably, NO_2_+NO_3_ were identified by the GLMs as being significantly affected by land use in the drainage basins.

**Figure 4.**
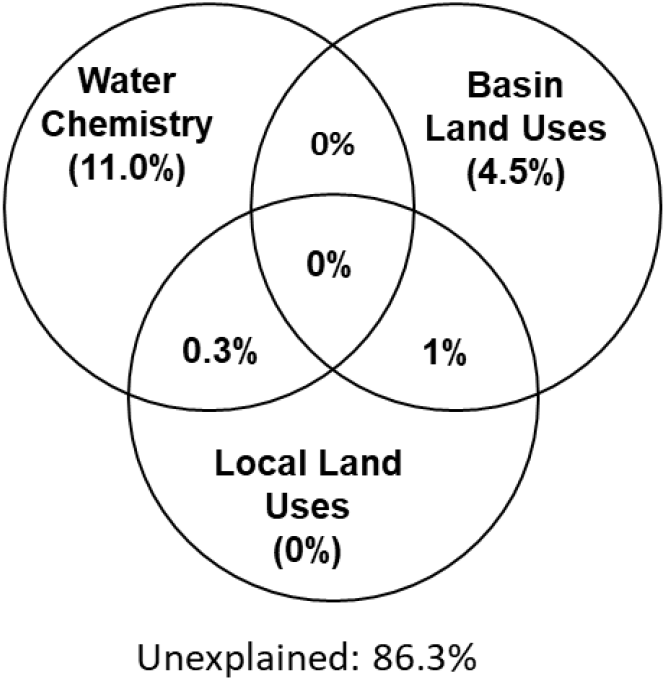
Water chemical properties represent the main explanatory variable for the microbiome composition (ca. 11% among all sites). Variation Partition Analysis performed with 3 data sets (basin and local land use as well as water chemistry) to explain the variation in the microbiome composition. The combination of land uses and water chemistry explain 13.7% of microbiome variability.

**Figure 5.**
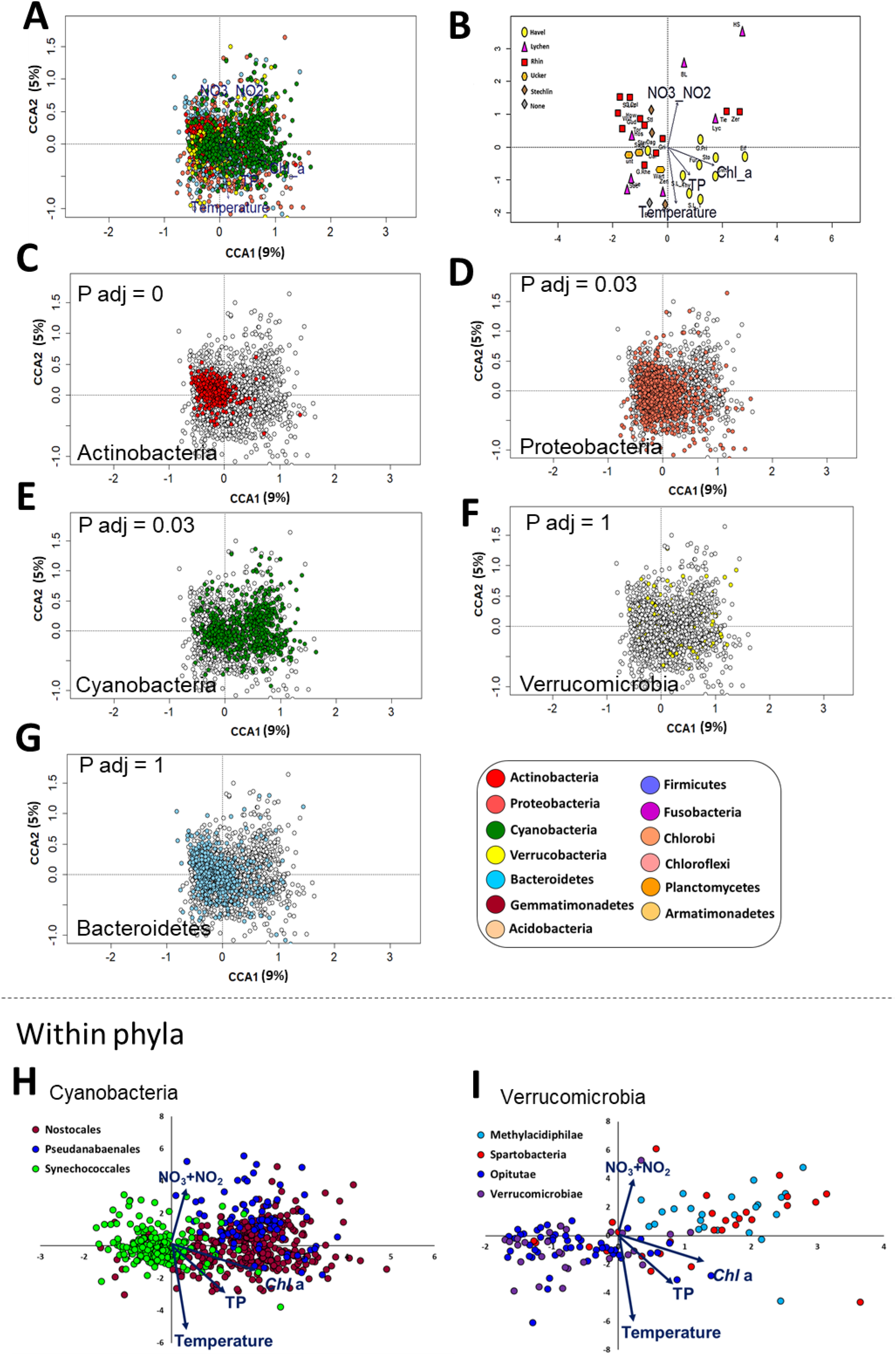
Differences between bacterial phyla in their association with different aquatic environmental conditions. Canonical Correspondence Analysis (CCA) plots are shown, with sampling locations and OTUs ordinated based on the four aquatic environmental parameters shown to significantly affect the microbiome composition. **A) An overview of the ordination, showing the 2845 most abundant OTUs**. The following panels each highlight a different aspect of this ordination. **B) The same ordination as in A, with the OTUs removed** to show the relative location of the eigenvectors for the aquatic environmental parameters and the ordination of the sampling sites. **C-G) The same plot as in A, with specific microbial phyla highlighted. H-I). Separation in the CCA ordination between selected lineages within the Cyanobacteria (h, orders) and Verrucomicrobia (I, classes)**. Note the clear separation between the Synechococcoales and the Pseudanabaenales and Nostocales along the first CCA axis with some differentiation between the latter two orders primarily along the second CCA axis (h). Within the Verrucomicrobia, the Methylacidiphilae, which are limited to the upper-right hand quadrant, possibly in association with high NO_2_+NO_3_ levels, are clearly separated from the other orders, except of the Spartobacteria.

### 3.3 Specific bacterial taxonomic groups are associated with sampling locations differing in their aquatic environmental conditions

Having identified TP, *Chl* a, NO_2_+NO_3_ and temperature as variables affecting the microbiome as a whole, we next asked whether we could identify specific microbial lineages affected by these environmental conditions. To answer this question, we analyzed the distribution patterns of OTUs belonging to different bacterial lineages along the CCA ordination axes. Most phyla were widely distributed in the CCA biplot (Figure 5), suggesting that, within each phylum, OTUs were affected differently by environmental factors driving the CCA ordination. However, Actinobacteria OTUs formed a tight cluster whose center was on the negative side of the x-axis (Figure 5C). This suggested that all OTUs in this group were affected similarly by the measured environmental variables. To statistically test this, we developed a permutation test to determine: a) whether OTUs in each lineage were clustered together more than expected (i.e. the distances between them were smaller than those between randomly selected OTUs. This suggests a coherent relationship with the factors constraining the CCA at the lineage level); b) whether the OTUs were associated (positively or negatively) with any of the eigenvectors of the specific aquatic parameters (i.e. the arrows in Figure 5B). We then applied these two tests to lineages at different phylogenetic resolution (phylum, class, order and family) to identify those that are coherently affected by the four environmental parameters tested (Supplementary Data File 2).

Of the five most abundant phyla, only Actinobacteria were not randomly distributed in the CCA plot, as determined by permutation analysis (Figure 5C-G, *P* < 0.05, Supplementary Data File 2). As a phylum, they were negatively associated with *Chl* a, temperature and total P. This suggests that, while the relative abundance of Actinobacteria was different between different sites, within this phylum OTUs behaved similarly, i.e. they are found at similar ratios at each location (Figure 2B). Proteobacteria showed a wider distribution range that was close to significance in our test (adjusted p-value = 0.066, Figure 5D). These results are in agreement with our observation that Actinobacteria and Proteobacteria OTUs dominate the core microbiome of all sites (Supplementary Figure 6A) and that within each of these phyla, a relatively large fraction of OTUs was core or cosmopolitan (Supplementary Figure 6B). In contrast, Verrucomicrobia, Cyanobacteria and Bacteroidetes did not cluster together in the CCA plot (although, as phyla, they were associated with some water parameters (Figure 5E-G, Supplementary Data File 2).

At a higher phylogenetic resolution, several lineages clustered in a non-random way and affected by the tested parameters (Supplementary Data File 2). In some cases, different lineages within the same phylum behaved similarly. For example, Acidimicrobiales and Actinomycetales orders, both belonging to the Actinobacteria, behaved similarly, being negatively associated with *Chl* a and total P concentrations, as well as with temperature. However, in other cases different lineages clearly clustered separately and were affected differently by the tested environmental factors, suggesting potential niche differentiation. Within the Cyanobacteria, the order Synechococcales was clearly separated in the CCA ordination from the Pseudoanabaenales and Nostocales (Figure 5H). The former was clustered non-randomly and was negatively associated with *Chl* a, NO_2_+NO_3_ and TP (Supplementary Data File 2), whereas the two latter orders were positively associated with these factors. Nostocales and Pseudanabaenales, in turn, differed in their correlation with temperature (positive and negative association with temperature, respectively). Notably, while Cyanobacteria appeared at all sampling sites (Figure 2B), few Cyanobacteria OTUs were part of the core microbiome (Supplementary Figure 6A), yet this phylum had the highest fraction of OTUs in the ‘’common’’ category among all analyzed phyla (Supplementary Figure 6B). This further suggests that different Cyanobacteria of different water bodies, potentially correlated with water quality and trophic status. Differences in the ordination of groups at the class level were also observed within the phylum Verrucomicrobia, with Opitutae clustered non-randomly and associated with low NO_2_+NO_3_ and *Chl* a, clearly separating from other classes such as Methylacidiphilae and Spartobacteria, both of which were positively associated with *Chl* a (although not clustered, Figure 6B).

**Figure 6.**
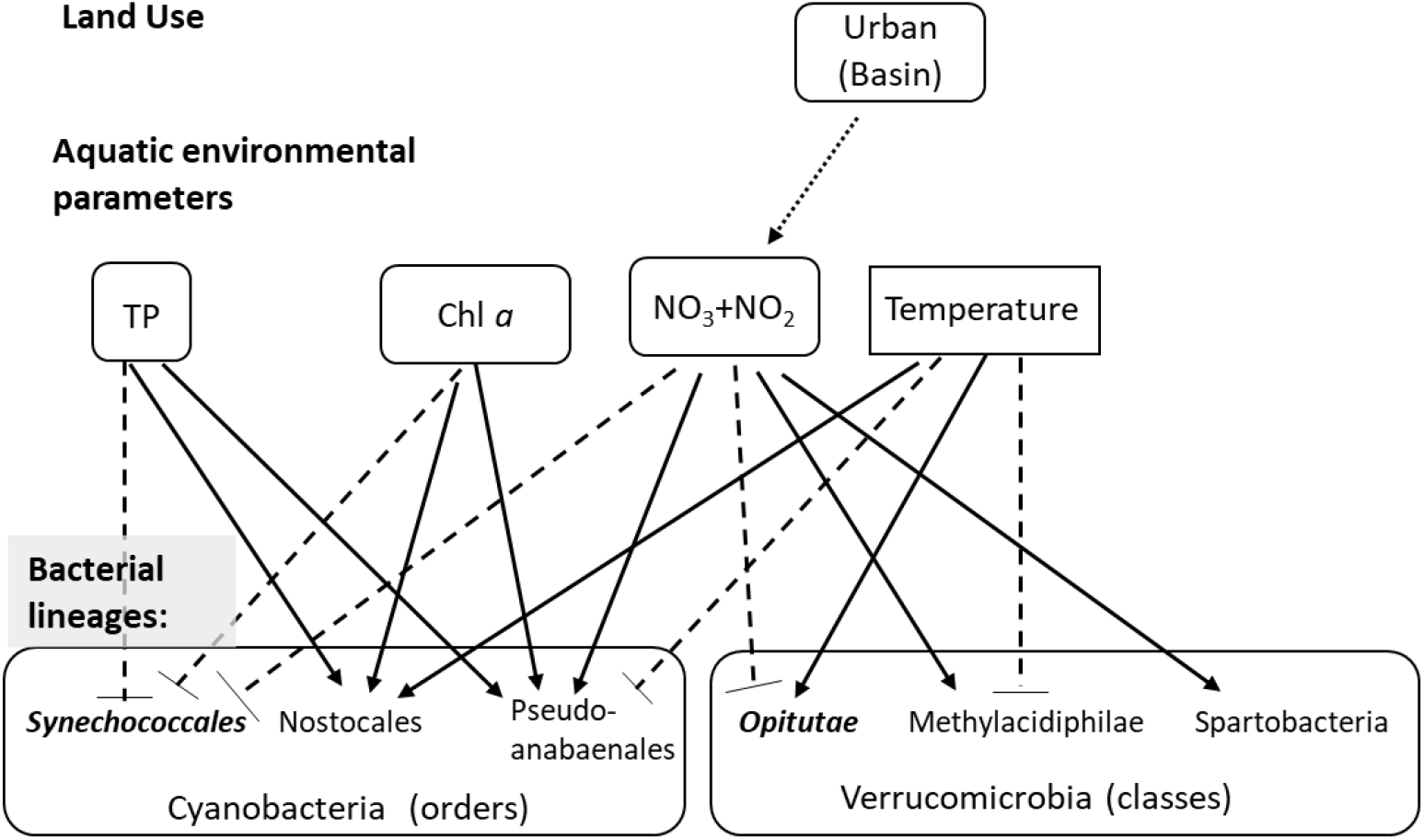
Conceptual model summarizing the observed effects of land use on water chemistry, and of water chemistry on selecetd microbial lineages. The first layer represents the effect of land use on aquatic environmental parameters (based on the GLMs), with the arrow showing the significant correlation between urban land and NO_2_+NO_3_. The seconds layer shows the effect of the four environmental parameters shown to sigificantly affect the microbiome structure (based on the CCA analysis) on selected microbial linages within the Cyanobacteria and Verrucomicrobia. Solid arrows show significant positive correlations (based on the permutation analysis), dotted blunt-ended lines show negative correlations. Lineages that are more clustered than randomly expected are in bold and italics.

## 4. Discussion

Quality and chemical properties of freshwater ecosystems, aquatic microbial populations, and human health are tightly interlinked. In this study, we analyzed the microbiomes of a network of partly-interconnected lakes, which belong to several different drainage basins. We aimed to identify to what extent land-use in the drainage basins of the lakes affects both environmental conditions in the water and lake microbiomes. Several previous studies have identified correlations of land-use to various parameters of water chemistry, as well as to the abundance of specific members of the microbial community (Savage, *et al*., 2010, Katsiapi, *et al*., 2012, Beaver, *et al*., 2014, Davis, *et al*., 2015). However, few studies have attempted to determine whether land use controls the microbiome structure as a whole (Glöckner, *et al*., 2000, Van Rossum, *et al*., 2015). Importantly, in our analyses, we used tools that are relatively accessible to water quality managers: land-use maps are available for many regions of the world, and are often used for land management using GIS (Joerin, *et al*., 2001, Zhang, *et al*., 2015). The water quality measurements performed using hand-held instruments, and measurements of inorganic nitrogen compounds and total phosphorus were performed using commonly-applicable and standardized methods (Grasshoff & Kremling, 1983, Grasshoff & Ehrhardt, 1999, Louca, *et al*., 2016). Our results suggest a conceptual model (schematically presented in Figure 6) whereby NO_2_+NO_3_ is related to a specific land use within the drainage basin of each water body (urban areas). In turn, NO_2_+NO_3_, together with total phosphorus *chl* a and temperature, are related to the relative abundance of specific microbial linages at different phylogenetic resolutions.

Previous studies have shown that specific environmental parameters affecting microbial community structure may vary between studies, freshwater systems and the time of sampling (e.g., Crump, *et al*., 2007). Temperature is often identified as an explanatory variable (Van der Gucht, *et al*., 2007), but this is not always the case (Savage, *et al*., 2010). *Chlorophyll* a and total P concentrations are also sometimes identified as important variables (Yannarell & Triplett, 2005), but not always (Crump, *et al*., 2007, Van der Gucht, *et al*., 2007, Niño-García, *et al*., 2016), and these two parameters may have a stronger effect on particle-associated rather than free-living bacteria (Savio, *et al*., 2015). In our dataset, *chl* a and total P concentrations (which are often used as quantitative measures of lake trophic state) were somewhat related to the relative abundance of Cyanobacteria (Supplementary Figure 2), especially Nostocales and Pseudanabaenales (Figure 5). While pH is often identified as important (Yannarell & Triplett, 2005, Crump, *et al*., 2007, Savio, *et al*., 2015, Niño-García, *et al*., 2016), it was not a statistically significant explanatory variable in our analysis. Finally, consistent with our results, NO_2_+NO_3_ have been shown to affect community structure in some cases (Kozak, *et al*., 2019), whereas in other studies total N (which was not measured here) was shown to be important (e.g., Van der Gucht, *et al*., 2007).

Below we first discuss specific links between land use types, water quality, and microbiome composition, as well as the association between specific lineages and water quality parameters. We then mention the caveats of our study and suggest ways to improve future studies.

### 4.1 What determines the microbiome - land use, mass transfer or biogeography?

Our initial hypothesis was that two types of factors would structure the microbial communities: i) environmental conditions within the water body, which are partly determined by land use within the drainage basin of each water body (often termed “species sorting”, e.g., (Van der Gucht, *et al*., 2007, Logue & Lindström, 2010), and ii) mass effects due to bacterial transport along the streams connecting the various sampled lakes. The relative importance of these two processes differs between different freshwater systems, with mass effects generally thought to be more prevalent in streams or smaller lakes with low water retention time and species sorting more prevalent in larger lakes with higher water retention times (Mašín, *et al*., 2003, Lindström & Bergström, 2004, Crump, *et al*., 2007, Logue & Lindström, 2010, Niño-García, *et al*., 2016). However, not all lake systems are alike, example distance-decay relationships (which suggest mass effects) were observed in some lake systems (e.g. (Van der Gucht, *et al*., 2007, Nelson, *et al*., 2009), but not in others (e.g. Niño-García, *et al*., 2016). Our results show no separation of bacterial communities by basin and no distance-decay patterns (Figure 2C, S2A). We note, however, that we did not measure the actual flux of water between the lakes, which could strongly affect the rate of emigration between the lakes (see “caveats”, below). Given the currently available data, it seems likely that, at the geographic scale and time-points we studied, there is no clear limitation on the dispersal of bacteria between the lakes. Rather, environmental conditions within each water body strongly affected the microbiome structure, and these were partly determined by land-use in the drainage area of each water body (Figure 4).

The effect of water quality on the microbiome was quantitative rather than qualitative, resulting in differences in the relative abundances of different microbial groups or OTUs rather than driving some of them locally extinct (Supplementary Figure 6). Indeed ~50% of the OTUs were found in >90% of the sampling locations, suggesting a “core” microbiome that is found at all sites, albeit with different relative abundancesn (Savio, *et al*., 2015). In organism-associated microbiomes, it has been suggested that microorganisms that are commonly found across locations or individuals (thus forming the core microbiome) may be critical to the function of the microbial community, and indeed may define a “healthy” community (Shade & Handelsman, 2011, Hernandez-Agreda, *et al*., 2017). Such studies are still rare for aquatic microbiomes, yet it is possible that better understanding of the dynamics of the core microbiome may help understand how microbial communities are related to – and affect – water quality.

Several studies have investigated the association between land use types and water properties (Quinn, *et al*., 1997, Larned, *et al*., 2004, Mehaffey, *et al*., 2005, Schoonover & Lockaby, 2006, Reimann, *et al*., 2009). These studies generally suggest higher dissolved nitrogen and phosphorus concentrations in waters surrounded by high coverage of urban areas and agriculture lands (Wang & Yin, 1997, Ghai, *et al*., 2014). In our study, urban land was an important predictor for (and positively correlated with) concentrations of NO_2_+NO_3_ (Figure 3B-C, Supplementary Tables 1, 2). The correlation between urban area (which includes also industry) and NO_2_+NO_3_ concentrations is in agreement with the previously mentioned studies, and may be driven by discharge of waste and rain run-off from sealed surfaces, both containing high nitrogen levels. Unlike previous studies, which observed a negative correlation between forested area and total phytoplankton as well as Cyanobacteria biomass (Catherine, *et al*., 2008), we did not observe such an effect, despite having sampled in lakes encompassing a wide array of trophic states. Neither did we observe an expected effect of agricultural land, e.g. through the effect of nutrient runoff (Carpenter, *et al*., 1998, Xu, *et al*., 2010), potentially because this region is characterized by low P fertilizer application and low livestock densities (Theobald, *et al*., 2016). Thus, in the region we studied, no clear correlation was observed between land use and total P or *chl* a concentrations (the low number of oligotrophic-mesotrophic lakes precluded an analysis of the trophic levels themselves). We note, however, that the interaction between land-use in lake catchments and the physics, chemistry and biology of the lakes is complex, with potentially conflicting drivers. For example, nutrients and dissolved organic matter exported from, e.g., forests into lakes may, on the one hand, promote phytoplankton and bacterioplankton growth, while at the same time increasing water color and subsequent light attenuation could result in changes in lake stratification (reviewed by Solomon, 2015).

While we were able to explain, to some extent, water parameters via land-use types, and the microbiome composition via water chemistry, we were unable to identify direct links between land use and the specific microbiome composition. An interpretation of these results might be that while land use in the drainage basin of each lake affects its microbiome indirectly, e.g., by impacting water chemistry, there is a relatively low direct effect of land use on the microbiome, e.g., by being a source for specific (presumably non-aquatic) microbes. In support of this interpretation, relatively few OTUs belonged to Acidobacteria (Supplementary Figure 6A) regarded as a representative phylum of soil bacteria (Janssen, 2006). Among these, only 0.04% of the total reads belonged to the *Solibacteres*, which have been considered as a marker for soil bacteria (Battistuzzi & Hedges, 2009). In a study of water bodies in the Ile-de-France region (Catherine, *et al*., 2008), land use was able to partly explain Cyanobacteria biomass, and in a subsequent study land use could predict to some extent the phytoplankton structure through the effect of land-use on water quality (Catherine, *et al*., 2016). Similarly, a land-use based model was able to partly predict the presence of genes involved in the biosynthesis of the Cyanobacteria toxin microcystin (Marmen, *et al*., 2016). In our dataset, even though the total fraction of Cyanobacteria is correlated with *Chl a* (R^2^ = 0.3 Supplementary Figure 3C), and the main groups of Cyanobacteria partition clearly into different niches based on the water chemistry (Figure 6A), we were unable to identify any land-use based model that could explain the variation in the relative abundance of these specific Cyanobacteria groups. Thus, while the potential of land-use to predict the abundance of Cyanobacteria and their toxins has been demonstrated, further work is needed to develop the appropriate methodologies and statistical tools that will identify linkages between land use types and microbiome structure. Finally, it is possible that land use might affect the function of the microbial population (at the genetic level) but that such an effect would not be clearly seen in the population structure, as measured by 16S amplicon sequencing (e.g., Louca, 2016).

Additionally, increasing the number of sampled locations will help to better make use of higher-resolution land-use maps, which may increase the ability to identify specific links between land-use and microbiome structure. Given the good accessibility of land-use data, which for much of the globe (including third world countries) can be derived from remote sensing, developing these methods should be of high priority for future research (Ndlela, *et al*., 2016).

### 4.2 Niche differentiation of freshwater bacterioplankton at the class and order levels

An important question in microbial ecology is whether there is a phylogenetic resolution at which environmental conditions can be best related to microbiome structure and function. Many studies focus on the phylum level (Lindström, *et al*., 2005, Briée, *et al*., 2007, Wolz, *et al*., 2018), since, at a broad scale, bacterial phyla differ in some basic functional traits including their ability to photosynthesize, or to utilize different carbon sources (Mitsui, *et al*., 1986, Camp, *et al*., 2009, Newton, *et al*., 2011). Indeed, a clear differentiation had been observed, in our dataset, between two groups of sites, those with a high relative abundance of Cyanobacteria, and those with a high relative abundance of Actinobacteria and Proteobacteria (Figure 2B). However, it has been suggested that this resolution is far too coarse to reliably infer community function, due to the high genomic and functional variation within phyla and even within individual bacterial lineages (Newton, *et al*., 2011, Singer, *et al*., 2016). Our results suggest that some bacterial lineages clearly partition between different environments already at low phylogenetic resolution, namely that of bacterial classes and orders. Several of these patterns were in agreement with our previous expectations based on literature data. For example, in freshwater environments, it is generally accepted that the relative contribution of Pico-Cyanobacteria (e.g., Synechococcales) to total phytoplankton decreases with higher trophic status. However, temporal patterns can be also important (Fukushima, *et al*., 2017). This may be attributed to the ability of Pico-Cyanobacteria to acquire major nutrients and trace metals at sub-micromolar concentrations, a capability that enables them to occupy relatively oligotrophic niches (Callieri, 2017). Larger phytoplankton (e.g., Nostocales) fills in niches with higher nutrient concentrations, and hence is physiologically adapted to proliferate at rich nutrient waters like fishponds and hypertrophic reservoirs (Cires & Ballot, 2016). The presence and density of Nostocales are of great interest from a water quality perspective due to their ability to produce a variety of secondary toxic metabolites (Sukenik, *et al*., 2012). In our samples, a clear separation could be observed in the CCA analyses between Synechococcales, associated with conditions of low *Chl a* and TP, and Nostocales, associated with the opposite conditions (Figure 5). Pseudanabaenales, which are filamentous Cyanobacteria whose cells are smaller than those of the Nostocales, were associated also with high NO_3_+NO_3_ concentrations. The physiology and ecology of Pseudanabaenales are relatively poorly known (Acinas, *et al*., 2008), yet these organisms are quite dominant in many of the lakes we sampled (Supplementary Figure 6), providing an incentive for further studies. The niche differentiation between Synechoccocales, Nostocales and Pseudanabaenales is the process that underlies the relative scarcity of ubiquitous Cyanobacteria OTUs, which instead dominate the “cosmopolitan” group (Supplementary Figure 6), and further highlight the potential importance of understanding the dynamics of “core” microbes in relation to water quality.

Potential niche differentiation was also observed within the phylum Verrocumicrobia, which is ubiquitous but relatively less studied in freshwater environments (Figure. 5I, Newton, *et al*., 2011). Some members of this phylum were found to be associated with high nutrient concentrations or algal blooms (Eiler & Bertilsson, 2004, Kolmonen, *et al*., 2004, Haukka, *et al*., 2006). In our study Methylacidiphilae (and possibly also Spartobacteria) clearly separated from the other three classes (Pedosphaerae, Opitutae and Verrucomicrobias), and may be associated with high NO_2_ + NO_3_ concentrations and low temperatures. At least some members of the Methylacidiphilae are methanotrophs (Camp, *et al*., 2009), and one bacterium from this class (*Methylacdiphilum fumariolicum*) is a diazotroph, able to actively fix atmospheric N2 (Khadem, *et al*., 2010). While most methanotrophic Verrucomicrobia have been isolated in extreme hot and acidic environments, the association of potential diazotrophs (Methylacidiphilae as well as Pseudanabaenales) with high NO_2_+NO_3_ in lakes is intriguing, and their potential contribution to high inorganic N concentrations is worth to perform additional studies.

The separation between different orders in the CCA plots suggests that, in freshwater bacteria, niche differentiation may be discernable at intermediate phylogenetic resolution - higher than that of phyla but lower than that of specific species or strains. We also searched for specific OTUs that were associated with aquatic environmental conditions or land use categories, but found very few results, and when correlations were observed, they were relatively weak (maximum Pearson R^2^ was 0.53, Supplementary Table 3). Two OTUs, one belonging to Phycisphaerales (phylum Planctomycetes) and one belonging to Saprospirales (phylum Bacteroidetes), were positively associated with NO_2_+NO_3_ concentrations. Phycisphaerales have been found in ANNAMOX processes (Hu, *et al*., 2015), and Saprospirales are associated with specific soil parameters (King, *et al*., 2010, Hargreaves, *et al*., 2015), yet, the role of these taxa in freshwater is mostly unclear. Thus, in our dataset, no specific OTU can serve as an indicator species for specific aquatic environmental parameters or land uses.

As the amount of genetic data (e.g. 16S rDNA surveys and metagenomes) from freshwater locations rapidly increases, it will be important to explore the best phylogenetic resolution at which to identify correlations between water quality and microbiome structure. Such microbiome data can then be linked to the relative abundance of specific microbial groups such as those involved in nutrient cycling, toxicity or pathogenesis, which in turn are important traits for water quality management.

### 4.3 Caveats and Limitations

While we could explain a part of the variability in the microbiome (a maximum of 13.7% in the VPA analysis, Figure. 4, similar to other studies, (Lear, *et al*., 2013), a large part of the variability remains unexplained. This variability may be due to other aquatic environmental parameters we did not measure, for example the quantity and composition of the dissolved organic carbon, which feeds heterotrophic bacteria and clearly has an effect on aquatic microbiomes (Davis, *et al*., 2015, Logue, *et al*., 2015, Wei, *et al*., 2016). Nor did we measure hydrological properties of the systems such as the lake depth, turbulence, stratification or retention times of the lakes, which can also strongly affect microbial population structure (Kopylov, *et al*., 2002, Nogueira, *et al*., 2002, Humayoun, *et al*., 2003, Carini, *et al*., 2005, Lindström, *et al*., 2005, Percent, *et al*., 2008). We also did not include in our analysis the potential contribution of groundwater, even though this might be significant. Lake retention times and potential groundwater inflow, for example, may affect the rate of population change and the relative impact of neutral processes such as emigration compared to effects of environmental drivers. While clearly of importance, often many of these hydrological parameters remain unknown, including for the majority of the lakes we studied, some which have been studied intensively for over 50 years (Allgaier & Grossart, 2006, Casper, 2012). More broadly, we include in our analysis only “bottom-up” controls, whereas biotic interactions such as grazing, phage infection and allelopathy can strongly impact population structure (Louca, *et al*., 2016).

Finally, our sampling approach, based on filtration on GF/F filters, assessed only part of the community, likely under-estimating the relative abundance of ultramicrobacteria (Proctor, *et al*., 2018). Many of these organisms belong to phyla that have yet to be cultured, and understanding their contribution to lake microbial dynamics represents a major future challenge. We also measured each location only once, relatively early during summer. Such a snap-shot does not represent the dynamic nature of the aquatic microbiome. For example, in several lakes that we studied, large Cyanobacteria blooms have been recorded, particularly during August (Mantzouki, *et al*., 2018). These blooms are often dominated by *Microcystis* species, yet these were relatively rare in our samples. Clearly, extended time-series are needed to gain a better understanding of the processes affecting aquatic microbiomes in relation to land use.

## Acknowledgements

We thank Elke Mach and Uta Mallok for chemical analyses, Liat Linde and Doron Fogel for the DNA extraction and the MIBI group for valuable discussions

This manuscript has been released as a Pre-Print at (Marmen *et al*., 2018).

## Author contributions

SM, HPG and DS designed study, SM, LG, HPG and DS collected samples, LB, LG, HPG and DS analyzed the geographic data, SM, LB, AA, AM, ML and DS analyzed the genetic data, SM and DS wrote the manuscript with contributions from all co-authors.

**<u>All authors have approved the final manuscript for publication</u>**

## Supplementary Material

### 1. Supplementary methods

#### 1.1 Study region and delineation of drainage basins

The sampling area lies within the region of Brandenburg, in northern Germany (Figure 1, Supplementary Figure 1). The area is relatively flat (18-89 meters above sea level) and contains hundreds of lakes, ponds and depressions that were formed by glacial action during the Pleistocene. Three main basins drain this region. From west to east, these are: 1) The Rhin basin, which flows south and then south-east, joining the Havel basin near Oranienburg; 2) The Havel basin itself, which flows generally south from the area of Lake Müritz, joining the Lychen sub-basin (which flows from the east) at Stolpsee (nearby Fürstenberg). It then continues south through Oranienburg to Berlin. The Havel then turns west and drains into the Elbe, itself draining into the North Sea, northwest of Hamburg; 3) the Ucker basin, flowing north through Oberuckersee and Unteruckersee and finally draining into the Szczecin Lagoon and the Baltic Sea. These three main basins can be subdivided into smaller basins.

The drainage basin of each lake was defined as the line connecting the highest points from which the surface water flow enters each of the lakes. Since the lakes are inter-connected, the drainage basin of each lake contains also those of the lakes upstream to it. Due to the flat topography of the region automatic basin detection using GIS was not satisfactory, and thus the basins depicted in Figure 1 and Supplementary Figure 1 were delineated manually using a combination of topographic maps produced from the Digital Elevation Model layer with a resolution of 0.0008333 decimal degree and high-resolution hiking and biking maps (Kompass Fahrradkarte, 1:70,000). Human structures such as roads were not informative due to the flat terrain. Literature surveys and site visits by the sampling team were used to clarify connectivity and flow conditions at specific locations. Notably, due to the flat terrain and the long history of water use for agriculture, aquaculture and transportation, the region includes also many artificial canals, weirs and ponds, often connecting between different basins. In order to simplify the analysis, the exchange between the basins was neglected. Additionally, Lake Müritz primarily drains north-east into Lake Kölpinsee and from there east into the Elde and ultimately into the Elbe. However, the lake also drains south at Rechlin into the Havel basin. The border between the Havel and Elde basins was delineated, based on the topography surrounding the lake, as shown in Figure 1.

Our 46 sampling sites varied between 7 different sub-basins as follow: 11 samples at Havel basin, 10 at Lychen, 12 at Rhin, 5 at Stechlin, 4 at Ucker, 1 sample from Tollensesee, 1 sample from Berlin Spree and 2 samples from lakes that are not part from any drainage basin (marked as “none” in Figure 1). Not all samples from the same basin are directly connected, for instance, water from the lakes in the Rhin’s northern part (12-15 in Supplementary Figure 1) meets water from the lakes in the Rhin’s eastern part (18, 19, 21-23) only downstream, near sites 16 and 17. Thus, in some cases sub-basins should be considered for the analyses (see main text and Sup Supplementary Figure 5).

The land uses, obtained from the Corine Land Cover map (CLC, http://www.eea.europa.eu/publications/COR0-landcover), were divided into five categories: Artificial areas, Agriculture (further divided for GLM analysis to heterogeneous agriculture, pasture lands and arable lands), Forests and natural areas, Wetlands and Water bodies (Figure 1). Sample locations were selected in order to represent a wide diversity of environmental parameters and land uses (Supplementary Data File 1, Supplementary Figure S1).

#### 1.2 Estimation of the trophic status of the lakes

The trophic classification of the lakes is based on measurements done or directed by the environmental agencies of the federal states Mecklenburg-Vorpommern and Brandenburg, following in general the rules of the water framework directive of the European Union. Such assessments are available for 25 out of the 46 lakes in the following link https://lfu.brandenburg.de/cms/detail.php/bb1.c.305410.de

Information for another 14 lakes was gathered from the following additional websites: https://www-docs.b-tu.de/fg-gewaesserschutz/public/projekte/uba_2/02_meck_pom.pdf https://www-docs.b-tu.de/fg-gewaesserschutz/public/projekte/uba_2/05_brandenburg.pdf https://www.lung.mv-regierung.de/dateien/glrp_ms_06_2011.pdf

Finally, the trophic state of the rest of the lakes was assessed based on the local expertise of the authors. Based on these assessments, there was one meso-oligotrophic lake, 8 mesotrophic lakes, 30 eutrophic lakes and 7 polytrophic lakes (including one fishpond, Supplementary excel file).

#### 1.3 DNA extraction, sequencing of 16S rRNA gene amplicons and sequence quality control

DNA extraction was performed at the Biomedical Core Facility of the Technion, Israel, using a combination of manual treatments and robotic extraction with the QiaCube robot. Briefly, filters were thawed, centrifuged for 10 min at 15000 × g and the preservation buffer was removed. Lysis buffer from the DNeasy Blood & Tissue Kit (Qiagen) was added and the samples were mechanically pounded with two 3mm sterile stainless steel beads at the speed of 30 Hz for 1.5 min using a TissueLyser LT (Qiagen). 30 μl of lysozyme were added, the tubes were incubated at 37°C for 30 min, followed by the addition of 25 mL proteinase K and 200□□l of the buffer AL with an additional incubation for 1 h at 56°C on a shaker. Finally, the tubes were centrifuged for 10 min at 5000 x *g* and the upper liquid was transferred to a new 2 mL Eppendorf tube for extraction by the QiaCube robot using the manufacturer’s instructions. The DNA quantity were assessed using Picogreen (Quant-iT PicoGreen dsDNA reagent, Invitrogen), and the quality (apparent size and degradation products) was assessed using a TapeStation (Agilent Technologies).

PCR reactions for amplifying the 16S rRNA gene fragments were performed in triplicate for each DNA sample using the following primers targeting the V3-V4 region: forward CS1_341F (5’-ACA CTG ACG ACA TGG TTC TAC ANN NNC CTA CGG GAG GCA GCA G-3’) and reverse CS2_806R (5’-TAC GGT AGC AGA GAC TTG GTC TGG ACT ACH VGG GTW TCT AAT-3’). The PCR reactions were performed in a volume of 25 μl, with 10 ng of DNA as PCR template. The PCR protocol was as follows: initial denaturation stage at 95°C for 5 min, followed by 28 cycles at 95 °C for 30s, 50°C for 30s, and 72°C for 60 s, with a final extension step at 72°C for 5 min. The enzyme used was BIOLINE 2x MyTaq Red Mix, and the PCR reaction was performed in a TProfessional Basic Gradient thermocycler (Biometra). Following the first PCR, triplicate samples were combined, and the products sent to the DNA Services Facility of the University of Illinois, Chicago, where a second PCR was performed to incorporate barcodes and sequencing adapters (CS1 and CS2). Sequencing was performed using a 2×250 base pair format on a MiSeq flow cell (V3 chemistry).

#### 1.4 16S analysis and statistical tests

Upon receiving the sequence data all sequences underwent quality control for Phix DNA removal using bowtie2 (Langmead & Salzberg, 2012), unmerged reads using PEAR (Zhang, *et al*., 2013) and incomplete, low quality scores and ambiguous bases sequences using MOTHUR Software V.1.36.1 using a phred score of 30, removing homopolymers longer than 8 nucleotides long and allowing no ambiguous bases (Schloss, *et al*., 2009). Following quality filtering, sequences were classified into different Operational Taxonomic Units (OTUs) based on 97% sequence similarity threshold using the “pick_de_novo_otus.py” command in Qiime (Caporaso, *et al*., 2010) and assigned taxonomy with the SILVA database version 128. Upon classification, an OTU table was generated and OTUs with fewer than 5 reads across the entire table were removed. At the end of analysis, the range of sequence number per sample was 35,775 – 75,704. This dataset, containing 45,240 OTUs, was used for all diversity analyses. For all multivariate statistical analyses and for the graphs presented in Figure 2B we used a smaller dataset which included only 3,611 OTUs with at least 50 reads across all sites. OTU numbers were log_10_ transformed. All calculations were performed in R software version 3.3.3 using the vegan package (Oksanen, *et al*., 2007).

Alpha diversity (Shannon index) was calculated using the ‘’diversity’’ function. The similarity in bacterial community structure among samples was calculated using a Bray– Curtis dissimilarity by ‘’vegdist’’ function. Gamma diversity was calculated in Excel by summing the number of sites where each OTU was present and dividing them into five categories: 1) Core OTUs, which were present at all the locations; 2) Cosmopolitan OTUs, which were present at more than 90% of the locations; 3) Common OTUS, present at more than 50% of the sites; 4) Rare OTUs, present in less than 10% of the sites; and 5) Unique OTUs, specific to only one site.

In order to explore whether samples from the same drainage basin have spatial similarity of bacterial communities, a non-metric multidimensional scaling analysis (nMDS) was carried out in R with ‘’metaMDS’’ function and a Bray-Curtis matrix. Sites were colored by basins.

Before performing multivariate analyses, we removed environmental parameters which were strongly correlated (Spearman *r* > 0.8, *P* < 0.05), with the exception of forests and arable lands, as the removal of one of these dominant land-use types significantly reduced the total fractional cover. Thus, the effect of these land-use types cannot be differentiated. The final environmental parameters selected were pH, temperature, ammonium, No_3_+NO_2_, total phosphorus and *Chl a*, generally referred as ‘’aquatic environmental parameters’’. Variation partitioning analysis (VPA, Legendre, *et al*., 2012) was used to describe the partitioning of variation in water microbiome among three data sets: ‘’Local Land Use’’ (the fractional land use within 500 m of the water body) and ‘’Basin Land Use’’ (the fractional land use within the drainage basin of each water body) and aquatic environmental parameters, see Supplementary Excel file 1, sheet “metadata”. For the analysis, we included thirteen land use types in the basin data-set and ten in the local (see Supplementary Data File 1). Land use types which were measurable at less in four sites were removed from analysis (e.g. Basin scale: Industrial areas, Construction areas, Artificial land cover, Crops, Vegetation, Open agriculture spaces and Wetlands; Local scale: Industrial areas, Artificial land cover, Herbaceous areas and Wetlands). We note that Forest and Arable lands have a strong negative correlation (Pearson *r* = −0.86), however, since in most basins these two land-use types contributes as much as 70-90% of the total area we did not remove one in order to avoid biases due to different total land-use values. The final table of land uses included six types in both scales: Urban fabric, Heterogenous agriculture, Pastures, Forests, Arable lands and Water bodies (see Supplementary Data file 1).

In order to validate the VPA results and further describe the effects of the different land use types on OTU patters, we performed canonical correspondence analysis (CCA) and permutation test (nperm=1000) using cca and anova.cca commands in R package ‘vegan’. We also considered whether the models have significant independent explanatory power by testing conditioned CCA. In the case of testing for model “Basin Land Use”, “Local Land Use” parameters were used as conditioning variables and vice versa.

VPA and CCA were also used to evaluate partitioning of variation in the bacterial community structure with three groups of explanatory variables: ‘’aquatic environmental parameters’’, ‘’Local Land Use’’, and ‘’Basin Land Use’’ (Supplementary Data file 1). Since varpart cannot be performed with missing data, we had to remove four sites (*i.e*. HAR, S.wen, Wen, F.P, Supplementary Data file 1) from the data-set before performing the analysis. Part of the removed samples are located around Berlin, for which we did not have detailed information on the drainage basin, and part were missing aquatic environmental measurements due to technical problems (I.e. Wan, Has and Hol).

**Supplementary Table 1:** Statistical values of each of the land uses in the GLMs correlations to water properties. The following abbreviations are used: U – Urban areas, Ar – Heterogenous agriculture, Pas – Pastures lands, Ag – Heterogeneous agriculture, For – Forests, W – Water bodies, *chl* a – *chlorophyll* a, TP – total phosphorus

| *Water parameter* | *Land uses* | *Level of land use* | *Model's $R^2$* | *Main contributing land uses for the correlation* |
| --- | --- | --- | --- | --- |
| $NO_2+NO_3$ | U+Ar+Pas+Ag+For+W | Local | 0.17 | Arable (+) |
|  | U+Ar+Pas+Ag+For+W | Basin | 0.74 | Urban (+) |
| *pH* | U+Ar+Pas+Ag+For+W | Local | 0.13 | – |
|  | U+Ar+Pas+Ag+For+W | Basin | 0.25 | Forests (-) |
| *Ammonium* | U+Ar+Pas+Ag+For+W | Local | 0.08 | – |
|  | U+Ar+Pas+Ag+For+W | Basin | 0.06 | – |
| *P* | U+Ar+Pas+Ag+For+W | Local | 0.02 | – |
|  | U+Ar+Pas+Ag+For+W | Basin | 0.14 | – |
| *Chl a* | U+Ar+Pas+Ag+For+W | Local | 0.06 | – |
|  | U+Ar+Pas+Ag+For+W | Basin | 0.078 | – |

**Supplementary Table 2.**
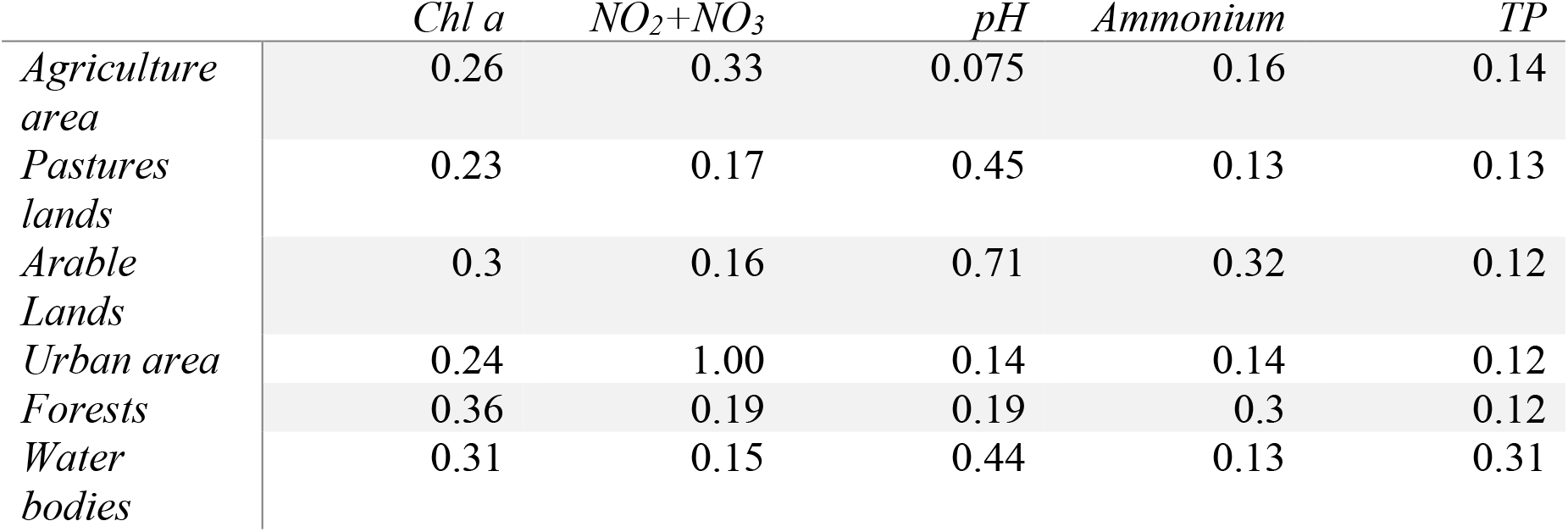
Importance values of each of the basin’s land uses in the GLMs. Abbreviations: *Chl a* – *chlorophyll a*, TP – total phosphorus

**Supplementary Table 3:** Spearman correlations between OTUs abundance and different environmental parameters. The only significant correlations occur between two OTUs and the water concentrations of NO_2_+NO_3_

| *Phylum* | *Order* | *Class* | *Family* | *Genus* | *Enviro-*<br>*nmental*<br>*parameter* | *r2* | *Adj*<br>*p val* |
| --- | --- | --- | --- | --- | --- | --- | --- |
| *Planctomycetes* | Phycisphaerae | Phycisphae-<br>-rales | NA | NA | NO<sub>2</sub>+NO<sub>3</sub> | 0.53 | 0 |
| *Bacteroidetes* | Saprospir-<br>ae | Saprospir-<br>-es | Saprospira-<br>-ceae | Aquirestis | NO<sub>2</sub>+NO<sub>3</sub> | 0.41 | 0 |

**Supplementary Table 4:**
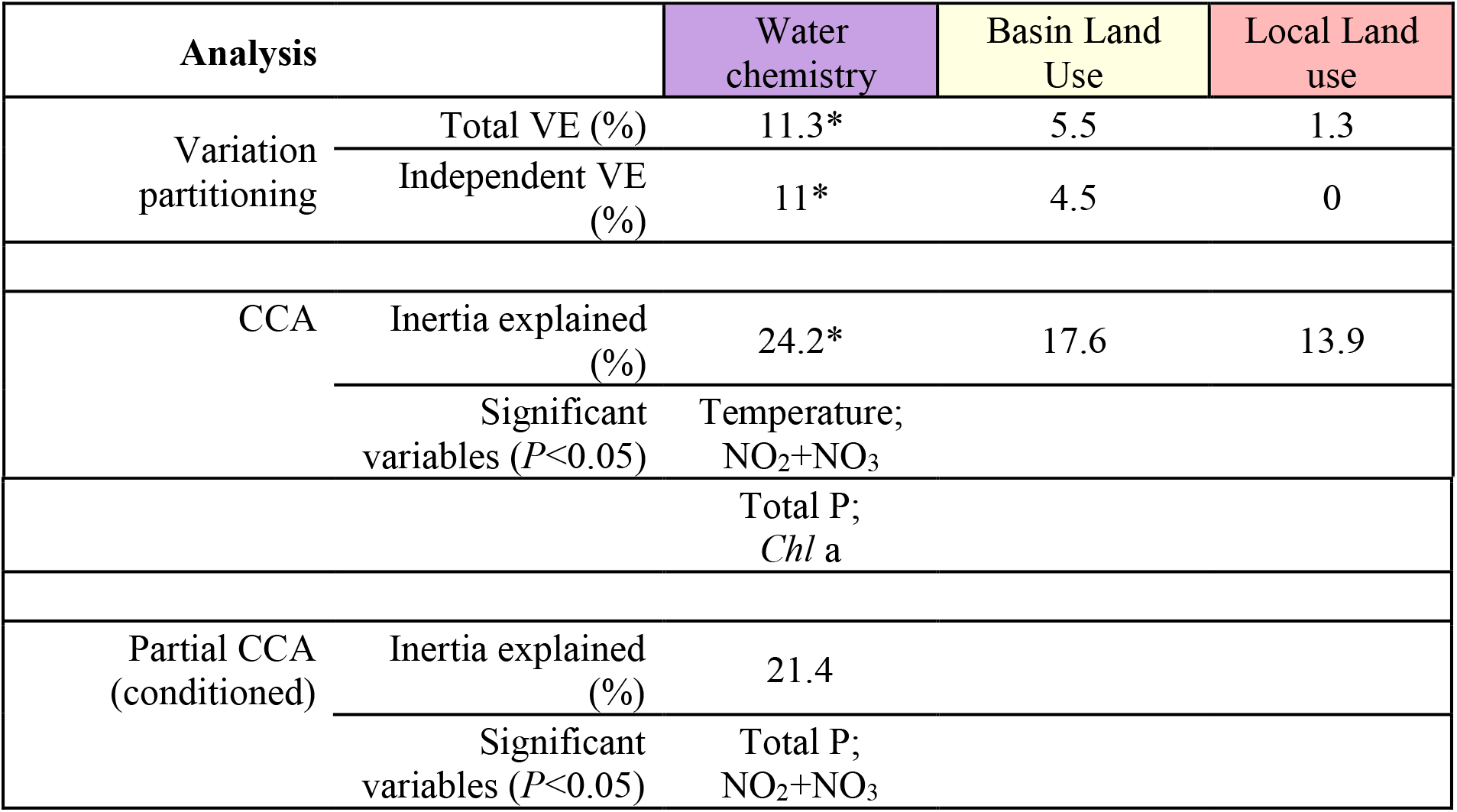
VPA and CCA results to describe the partitioning of water microbiome by variation in aquatic environmental parameters and land uses. Significance of the CCA models tested using Anova. * represents statistical significance.

## Supplementery Figures

**Fig S1.**
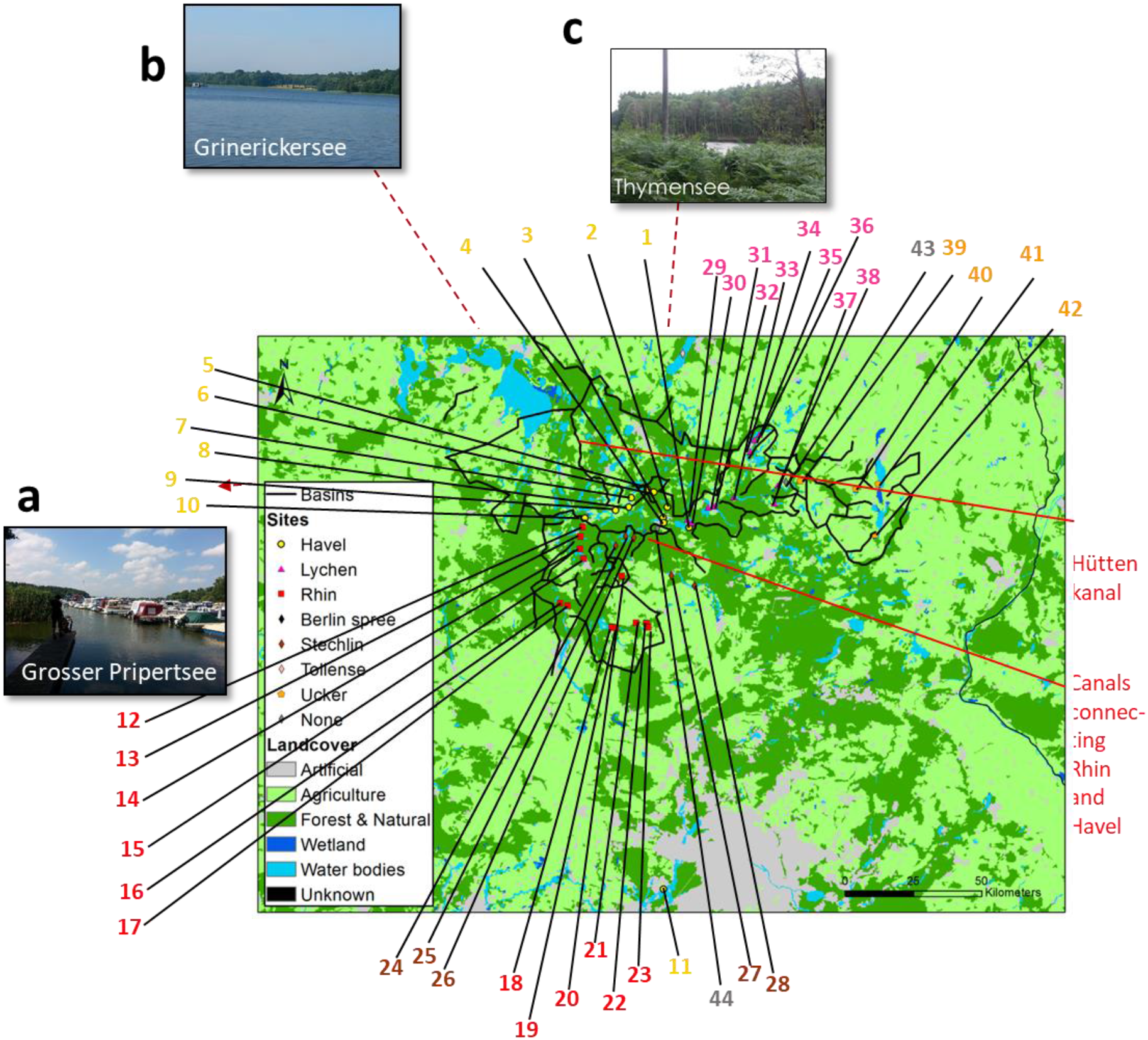
Detailed map of the sampling location, with the different sampling sites noted. Sampling sites were chosen to represent a high diversity in environmental gradients, e.g. variety in land use and water chemistry. These differences between the sites can be visualized - even before our measurement: **a-c)** Lakes of the Havel basin. **a)** Grosser Priepertsee is lake located within a town called Priepert and is used for boating and recreation, while **b)** Grienericksee is located next to the city of Rheinsberg with clear blue water. **c)** Thymensee is located in the middle of a forest area far away from any town. Arrows with numbers indicate sites as are presented in Supplementary Excel table. Red lines represent connectivity between drainage basins, as mentioned in the text. Notice that two of the sites studied (Hölzerner See and Tollensesee) are outside the scope of this map. Their exact coordinates can be found in the Supplementary Excel table under the numbers 45 and 46.

**Fig S2.**
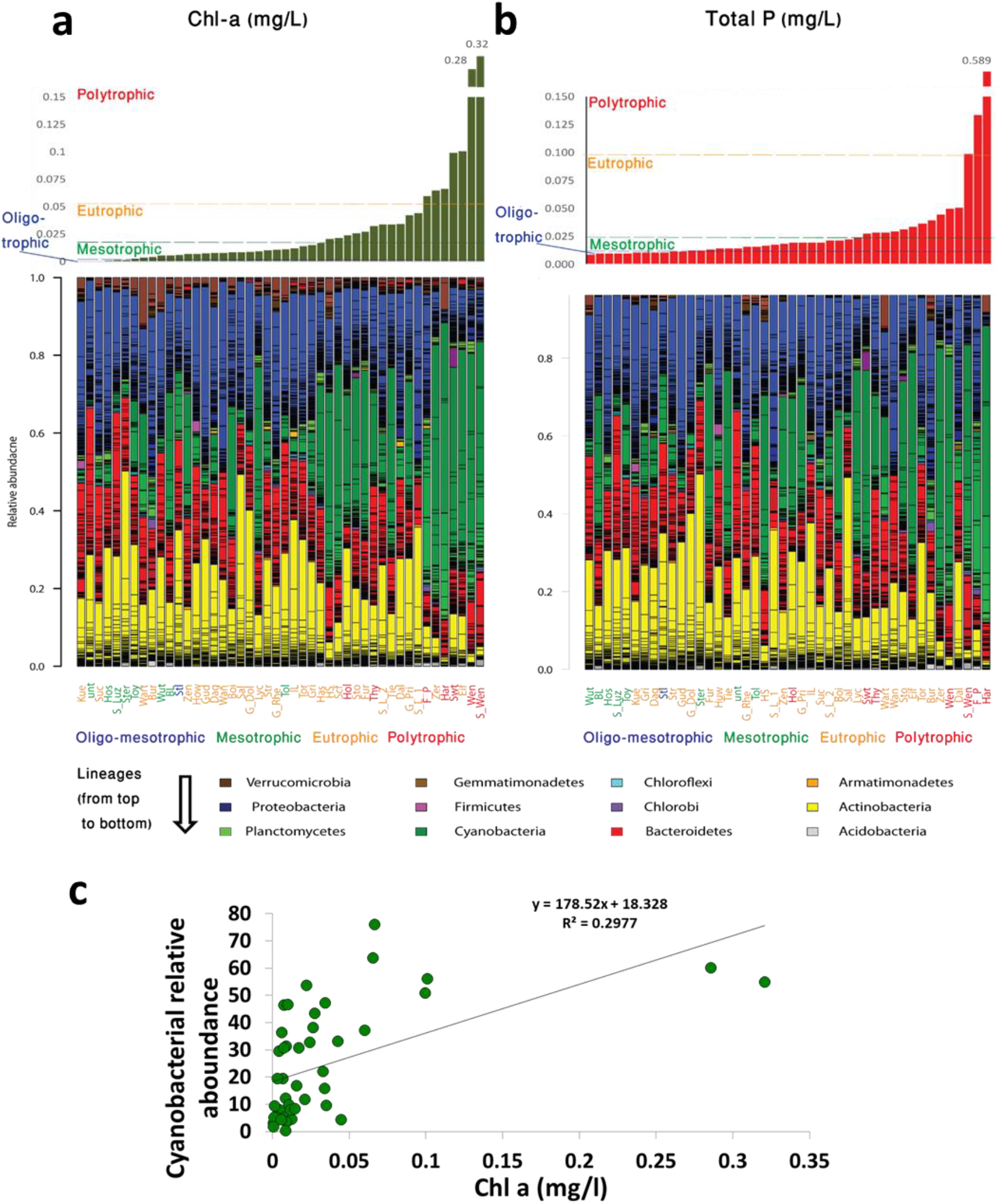
Relationship between trophic state and microbial community composition. a, b) The population structure of the samples lakes, ordered by chlorophyll (a) and total P concentrations (b) at the time of sampling. The codes for the location sites are colored based on the trophic state shown in the supplementary excel file, which is based on the monitoring programs of the German federal states Mecklenburg-Vorpommern and Brandenburg, following in general the rules of the water framework directive of the European Union (see supplementary text for more information). A slight tendency is seen towards increased relative abundance of cyanobacteria when *Chl* a (A) and total P (B) are high. C) **Weak correlation between chlorophyll concentration and relative abundance of cyanobacteria**. See also the NMDS of figure S3.

**Fig S3.**
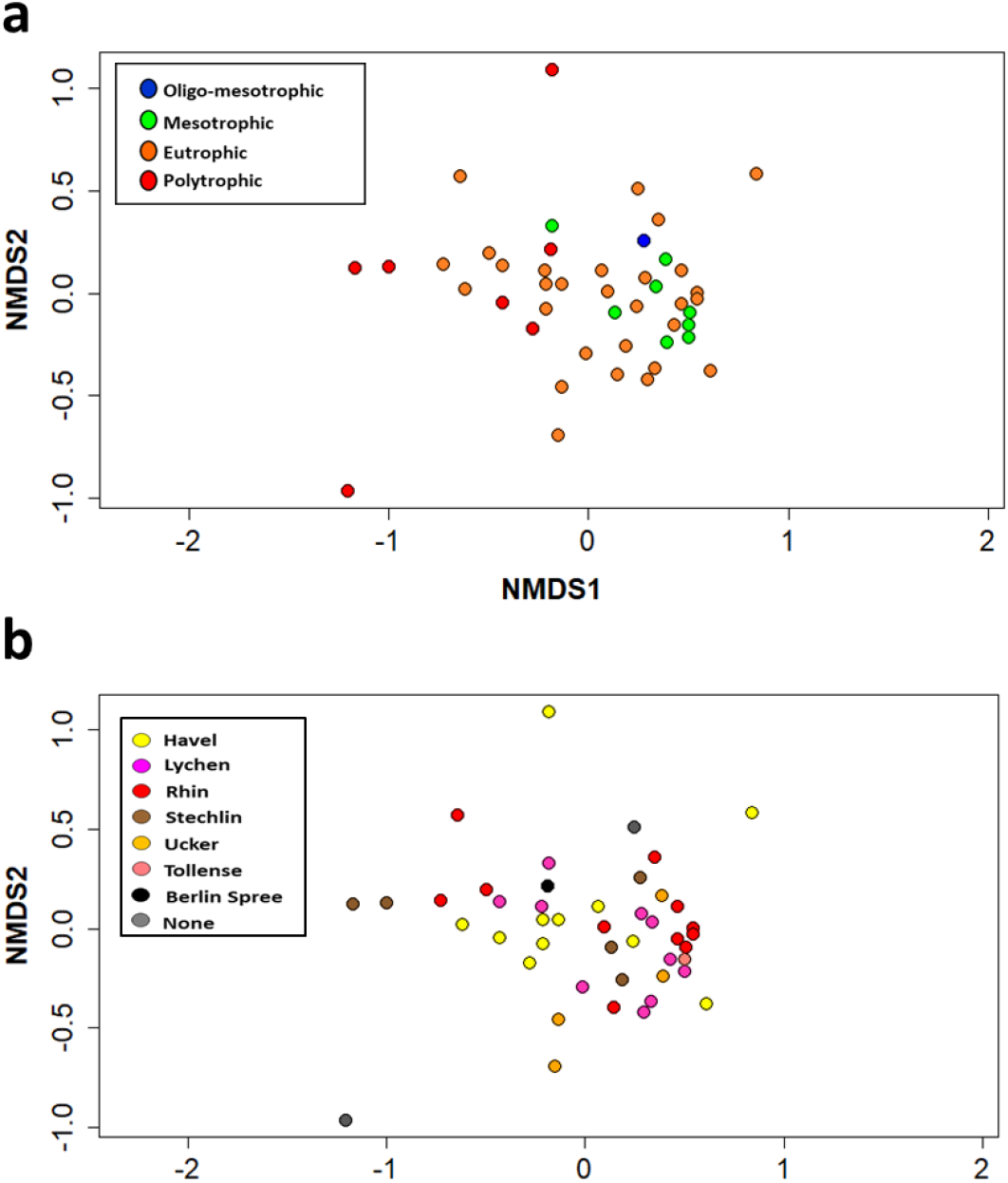
NMDS analysis of the bacterial population structure, colored by trophic level (a) and by drainage basin (b). The NMDS was calculated using Bray-Curtis dissimilarities. K=2, Stress= 0.16

**Fig S4.**
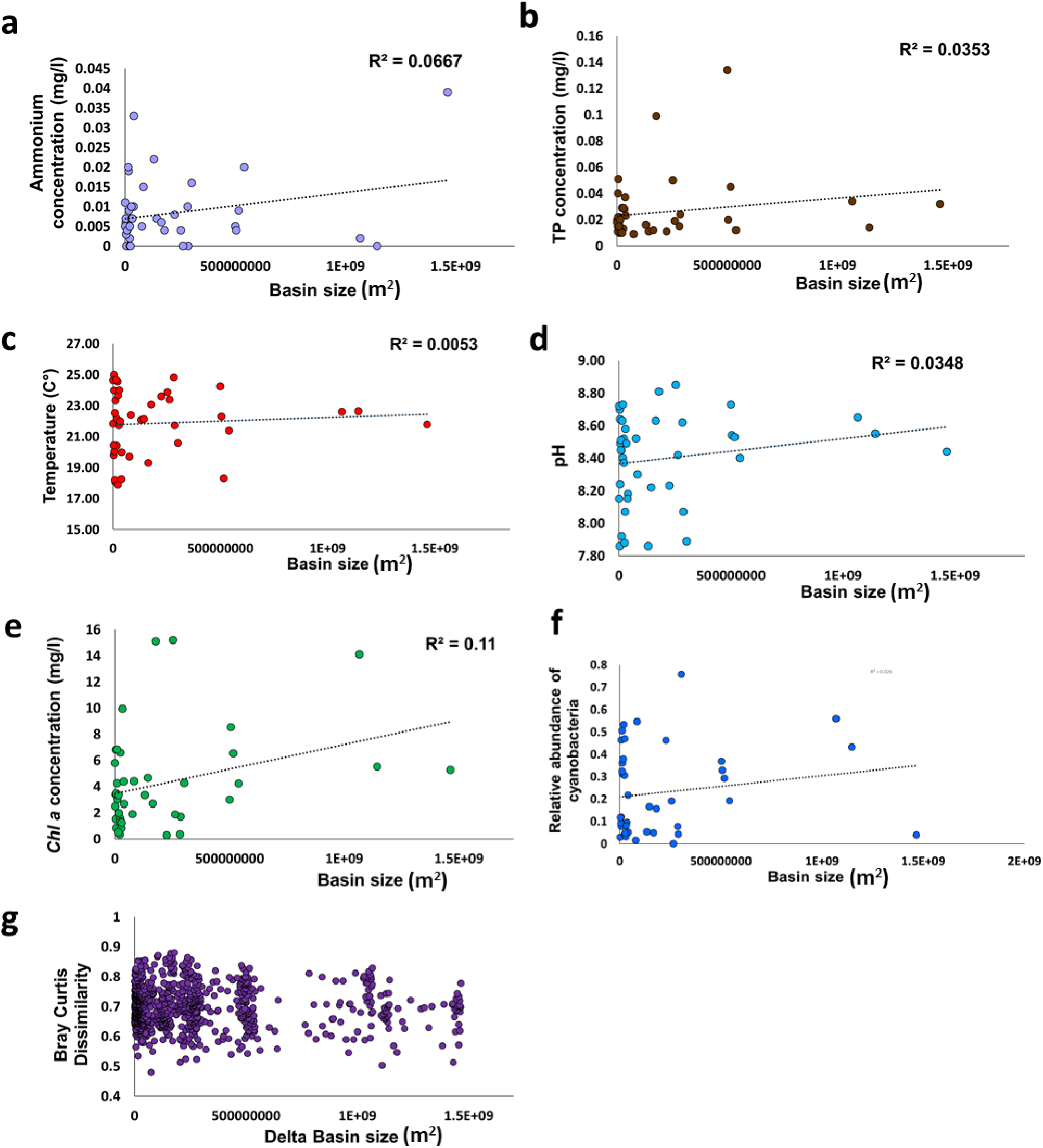
No clear correlation between basin size, water chemistry or microbiome composition. a-d) Drainage basin size vs ammonium (a), total phosphorus (b), temperature (c) and pH (d); no clear correlation can be seen. e) Low correlation between basin size and Chl *a* concentration, and f) No clear correlation between the fraction of cyanobacteria from the total microbiome and the basin size. g) No correlation between basin size and microbiome similarity. Each point represents the Bray-Curtis dissimilarity between two sampling points, plotted against the difference in size between the drainage basins of the two locations.

**Fig S5.**
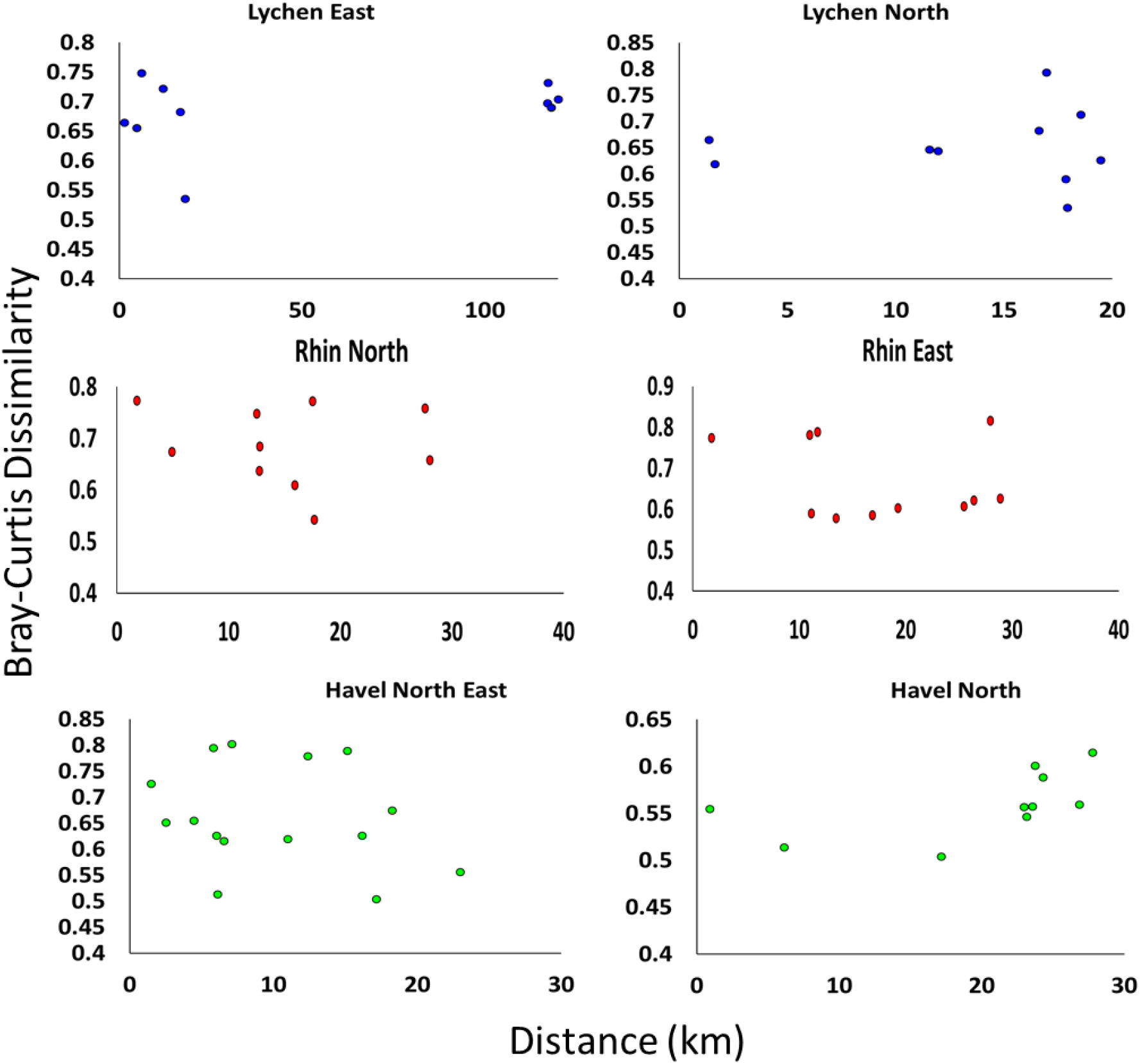
No distance decay patterns observed in sub-basins. The three large basins in the analysis were each divided into two smaller sub-basins, where all of the water bodies are directly connected. No distance-decay patterns were observed (Spearmans r ranged between 0.1 and 0.26, p values 0.3-0.7). The water bodies used for each sub-basin are as follows (see Supplementary excel File for water body codes). **Lychen East**: Lyc, Zen, Kue, Hos, Has; **Lychen north**: Lyc, Zen, HS, BL, Has; **Rhin North**: G.Rhe, Gri, Tie, Tor, Zer; **Rhin East**: Tor, Gud, G.Dol, Huw, Wut, Zer; **Havel North-east**: Thy, S.L_1, S.L_2, Sto, Fur, Swt; **Havel North**: G.Pri, Sto, EIF, Fur, .Swt

**Fig S6.**
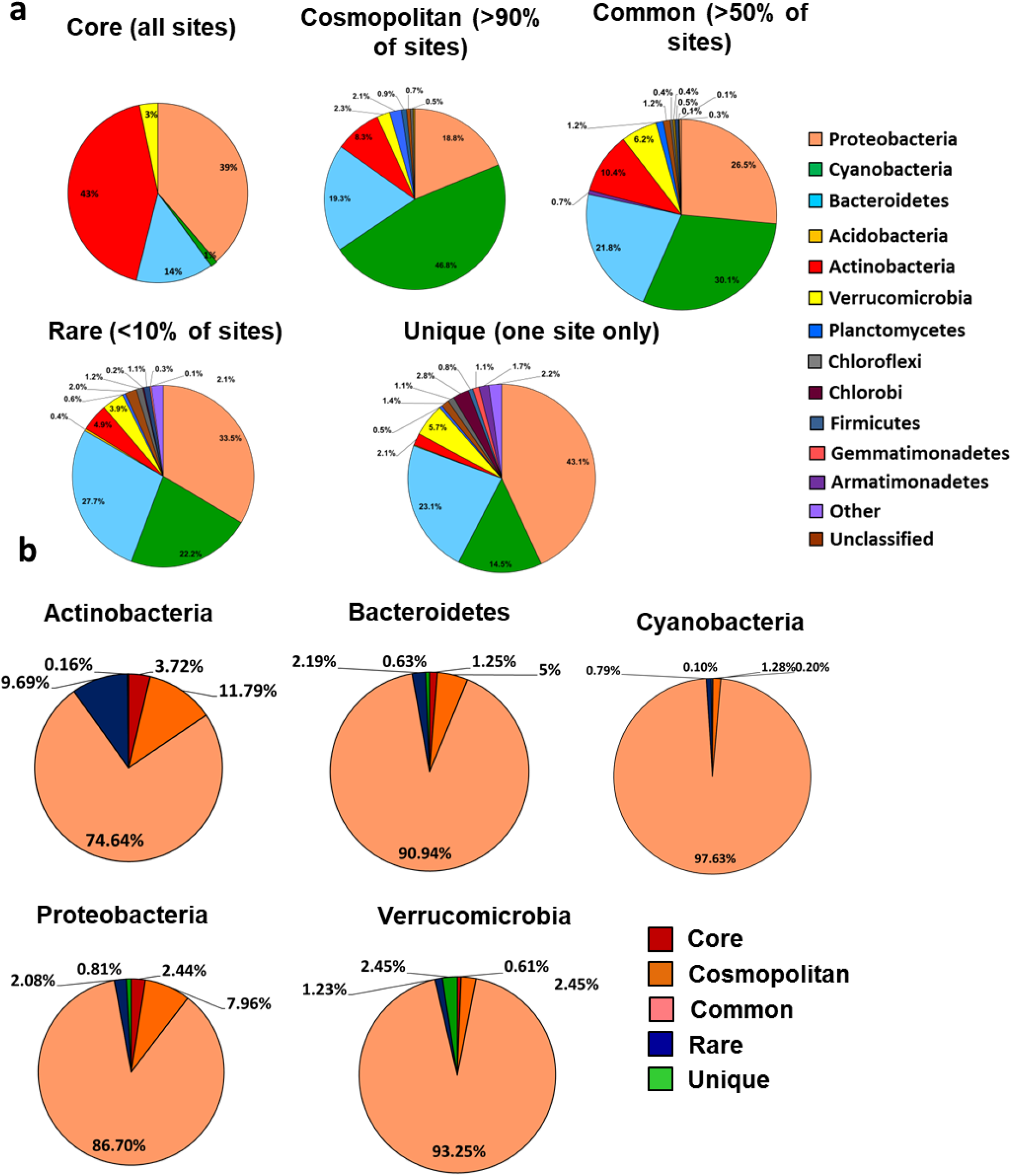
Relationships between different phyla and their relative abundance in the combined microbiome of all sampling sites. **a) Relative abundance of bacterial phyla among core, cosmopolitan, common, rare and unique OTUs**. The core OTUs, found in all sampling locations, are dominated by Proteobacteria and Actinobacteria. In contrast, few cyanobacterial OTUs are found in all sampling locations, whereas many are cosmopolitan (found in >90% of the locations**). B) Distribution of core, cosmopolitan, common, rare and unique OTUs among the different phyla**. Note the difference in the relative number of core and cosmopolitan OTUs between Actinobacteria, suggested to have similar communities in all sampling sites, and Cyanobacteria, which reveal niche differences between different orders.

**Fig S7.**
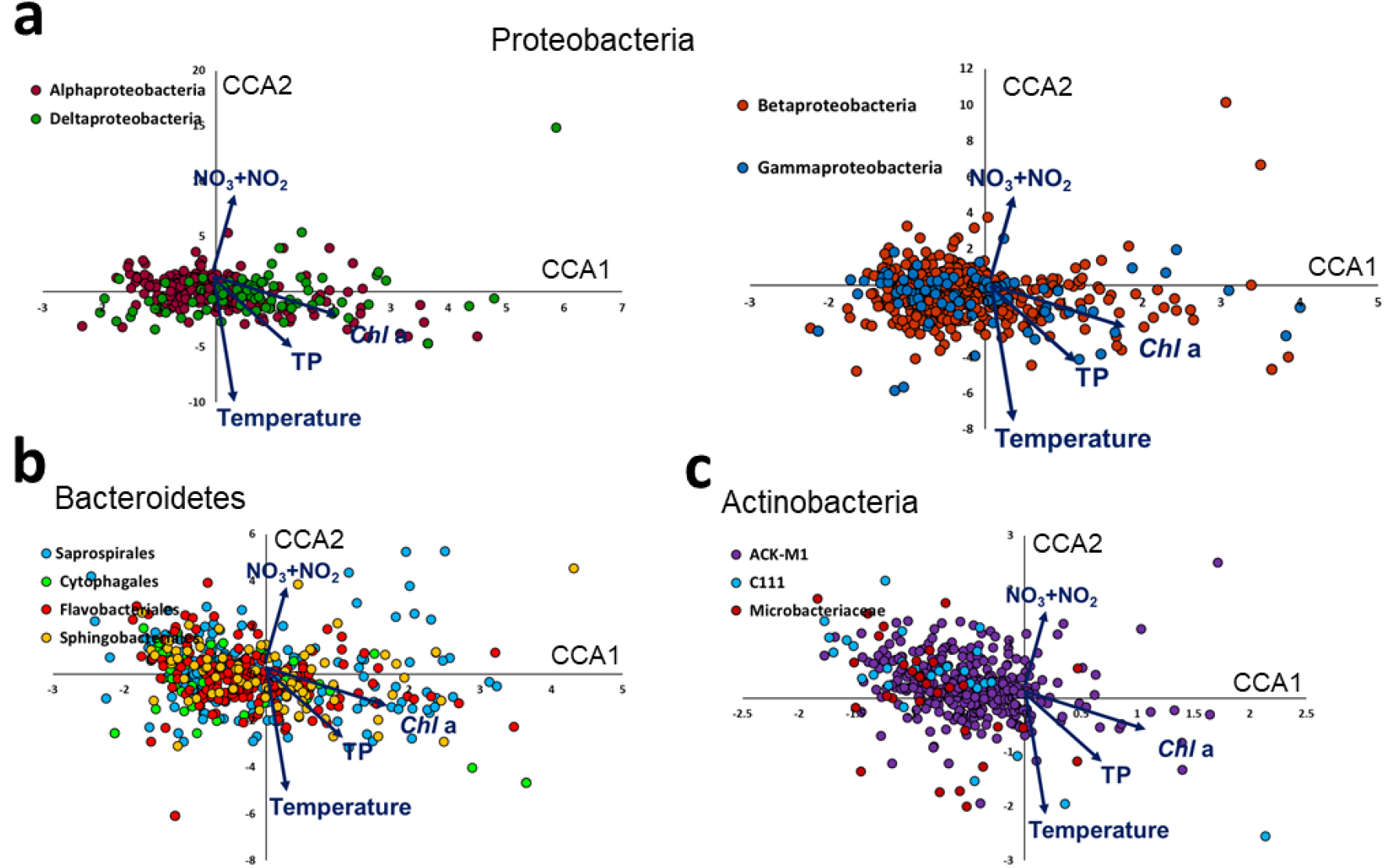
Absences of separation in the CCA ordination between several bacterial lineages: a) Proteobacteria orders, b) Bacteroidetes orders, c) Actinobacteria families. No clear differentiation is seen between the orders or families.

